# mAbClust with AlphaFold 3 avoids hallucinations to define a quaternary broadly neutralizing HCV epitope

**DOI:** 10.64898/2026.01.06.698032

**Authors:** Harry T. Paul, Jessica L. Mimms, Hanna Aboueid, Laura Sanchez Buitrago, Liudmila Kulakova, Xander Wilcox, Katrina Kuhn, Hao Zhang, Eric A. Toth, Nicole Skinner, Andrew I. Flyak, Gilad Ofek, H. Benjamin Larman, Thomas R. Fuerst, Stuart C. Ray, Andrea L. Cox, Justin R. Bailey

**Affiliations:** Division of Infectious Diseases, Department of Medicine, Johns Hopkins University School of Medicine, Baltimore, MD. USA; Institute for Bioscience and Biotechnology Research, University of Maryland, Rockville, MD, USA; Department of Microbiology and Immunology, Cornell University, Ithaca, NY, USA; Department of Molecular Microbiology and Immunology, Johns Hopkins University Bloomberg School of Public Health, Baltimore, MD, USA; Department of Cell Biology and Molecular Genetics, University of Maryland, College Park, MD, 20742, USA; The Abigail Wexner Research Institute at Nationwide Children’s Hospital, Columbus, OH 43205, USA; Department of Internal Medicine, College of Medicine, The Ohio State University, Columbus, OH 43210, USA; Institute for Cell Engineering, Division of Immunology, Department of Pathology, Johns Hopkins School of Medicine, Baltimore, MD, USA; Department of Oncology, Johns Hopkins University School of Medicine, Baltimore, MD, USA

**Author notes:** Lead contact: Justin R. Bailey 855 N. Wolfe Street, Rangos Building, Suite 520 Baltimore, MD 21205, USA.

## Abstract

The hepatitis C virus (HCV) envelope glycoprotein E1E2 heterodimer is the target of broadly neutralizing antibodies (bNAbs). Although prior studies have indicated that E1-dependent bNAbs are associated with spontaneous clearance of HCV, all E1-dependent human monoclonal antibodies (mAbs) have been isolated from individuals with chronic HCV infection. Here, we isolated E1-dependent bNAbs from an individual with high neutralizing antibody breadth who spontaneously cleared HCV, showing that these bNAbs bind to four distinct sites on E1E2. We also developed mAbClust, an algorithm that improves identification of accurate AlphaFold 3 (AF3) structure predictions of antigen-antibody complexes. We used AF3 and mAbClust to generate a high-confidence predicted structure of an E1-dependent bNAb in complex with E1E2, showing that this bNAb binds to a quaternary epitope spanning E1 and E2. This study identifies four neutralizing sites and a quaternary bNAb epitope associated with HCV control, which can guide HCV vaccine design. AF3 with mAbClust could have broad applications for accurate epitope mapping of antibodies.

## Introduction

There are currently about 50 million people infected with hepatitis C virus (HCV), with about 1 million new infections per year^1^. Incident HCV infections are outpacing the rate at which existing infections are being cured with direct acting antivirals (DAAs), further exacerbating the global health burden^1^. Along with increased access to treatment, a prophylactic vaccine against HCV is needed to achieve the World Health Organization goal for HCV elimination^1^. No protective vaccine currently exists, but there is cause for optimism, since about 25% of infected individuals spontaneous clear their infection without treatment, and these individuals then have roughly an 80% chance of clearing any subsequent HCV infection^2–6^.

Multiple studies of plasma antibodies^7–9^ and monoclonal antibodies (mAbs)^10–12^ have identified potent broadly neutralizing antibodies (bNAbs) against HCV, which can block infection with diverse viral strains in vitro and in animal models of infection^3,6,7,13–21^. These bNAbs target the envelope glycoproteins, E1 and E2, which form a heterodimer on the surface of virions that mediates interaction with cell surface receptors, viral fusion, and entry. Given the genetic and antigenic diversity of E1E2^22,23^, and the role of this variability in facilitating escape from antibody pressure^24–26^, understanding the landscape of E1E2 epitopes targeted by bNAbs is critical to vaccine design.

Many bNAb epitopes in E2 have been characterized^14,27–30^. This includes epitopes in the front layer, back layer, and beta sandwich of E2, targeted by bNAbs from individuals with chronic HCV infection as well as people who spontaneously cleared HCV without treatment^13,27,31^. In contrast, although analyses of polyclonal plasma antibodies have indicated that E1-dependent bNAbs are associated with spontaneous control of HCV^2,18^, very few E1-dependent human mAbs have been isolated, and all were isolated from individuals with chronic HCV infection. E1-dependent antibodies that have been characterized fall into two main groups. The first group binds E1 directly, like closely related mAbs IGH505/IGH520/IGH526^32^ as well as mAb H-111^33^. The second group comprises mAbs like AR4A^34^, which has binding residues entirely in E2^35^, but is E1-dependent because it binds only when E2 is part of a conformationally intact E1E2 heterodimer. To date, no E1-dependent human mAbs have been described that bind to quaternary epitopes spanning E1 and E2.

In addition to a lack of E1-dependent bNAbs isolated from persons with HCV clearance, another major limitation in understanding the role of these bNAbs in HCV control has been difficulty in obtaining structures of E1-dependent bNAbs in complex with E1E2, largely due to difficulty in expressing conformationally-intact E1E2 heterodimers. Only recently have investigators used cryogenic electron microscopy (cryo-EM) to generate the first structures of AR4A and IGH505 in complex with E1E2 heterodimers^35,36^.

The development of AlphaFold 3 (AF3) presents a new opportunity to map the binding sites of HCV bNAbs. AF3 is a generative artificial intelligence (AI) that predicts the molecular structures of complexes of multiple proteins^37^. AF3 could be particularly useful for understanding neutralization by E1-dependent bNAbs since determining structures experimentally has been particularly challenging. However, AF3 has not been tested with HCV-specific mAbs. In addition, although AF3 has shown great promise for accurately predicting structures of antibody-antigen complexes, a limitation in its use has been difficulty in distinguishing accurate AF3 structure predictions from inaccurate predictions^38–40^.

In this study, we developed a new method for isolation of E1-dependent bNAbs, which allowed us to examine the E1-dependent antibody repertoire in an individual with high neutralizing antibody breadth who spontaneously cleared HCV without treatment. We also developed and validated mAbClust, a new algorithm that improves identification of accurate AF3 predictions of antigen-antibody complexes. We then used AF3 with mAbClust to generate a high-confidence predicted structure of a novel E1-dependent bNAb in complex with E1E2, showing that this bNAb binds to a quaternary epitope with contact residues on both E1 and E2.

## Results

### Isolation of genetically diverse E1-dependent monoclonal antibodies (mAbs) from an Elite Neutralizer of hepatitis C virus (HCV)

We previously screened neutralizing breadth of early-infection plasma samples from 63 participants in the Baltimore Before and After Acute Study of Hepatitis (BBAASH), a prospective, longitudinal cohort of people who inject drugs with incident HCV infection^27,41^. We measured neutralizing breadth using a panel of genetically and antigenically diverse HCV pseudoparticles (HCVpp)^23^, designating the 10% of individuals in the cohort with greatest neutralizing breadth as Elite Neutralizers (EN)^27,42^ (Fig. 1A). One EN, designated C117, is of particular interest because their plasma neutralized 14 of 15 HCVpp (Fig. S1), and they naturally cleared their gt1a HCV infection without treatment and without detectable T cell responses^19^. We previously isolated multiple broadly neutralizing mAbs (bNAbs) targeting the E2 envelope glycoprotein from this individual using hybridoma technology^13^, but, due to technical limitations, had not investigated the repertoire of antibodies targeting E1-dependent epitopes.

**Fig 1.**
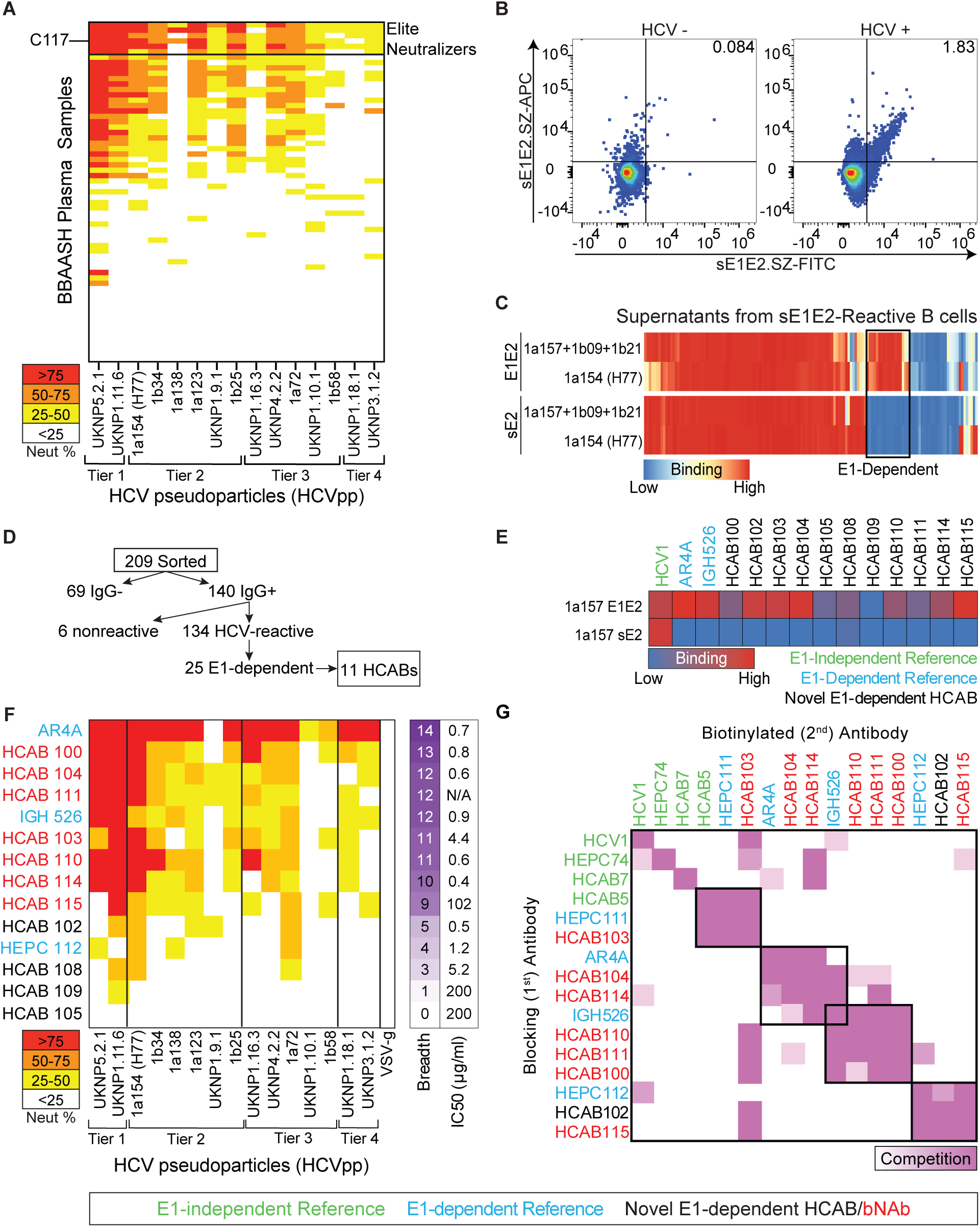
Isolation and characterization of genetically diverse E1-dependent mAbs from an Elite Neutralizer of HCV. (A) Neutralizing breadth of early-infection plasma samples from 63 participants in the Baltimore Before and After Acute Study of Hepatitis (BBAASH) cohort measured with a panel of antigenically diverse HCVpp with four tiers of increasing neutralization resistance. Participants are arranged from greatest to least neutralizing breadth, with a horizontal line demarcating the top 10%, designated Elite Neutralizers. Participant C117 is an Elite Neutralizer who spontaneously cleared their infection with neutralization of 14 of 15 HCVpp. Heatmap represents % neutralization values measured with 1:100 diluted plasma. Values are the average of two independent experiments, performed in duplicate. (B) Dual-fluorophore sE1E2.SZ staining of class-switched B cells shows an E1E2-reactive population in an HCV+ individual compared to an uninfected control. (C) Binding of IgG in single B cell culture supernatants to soluble E2 (sE2) or lysates of cells expressing full-length E1E2. Proteins were derived from strain 1a154 (H77) or a mixture of strains 1a157, 1b09, and 1b21. Single E1E2-reactive B cells from C117 were flow-sorted and cultured to stimulate antibody production. 25 of 134 HCV-reactive supernatants (19%) bound E1E2 and not sE2. Heatmap represents Z-score of OD_450_ binding values by ELISA of supernatants. Values were measured in a single experiment. (D) Yield at each stage of antibody isolation, leading to 11 E1-dependent recombinant monoclonal antibodies (mAbs), designated Hepatitis C Antibodies (HCABs). (E) Binding of HCABs to sE2 or E1E2 lysates derived from strain 1a157. E1-independent reference mAb HCV1 (green) and E1-dependent reference mAbs AR4A and IGH526 (blue) are shown as controls. Heatmap represents OD_450_ binding values by ELISA of mAbs at 100 µg/mL. Values were measured in a single experiment, performed in duplicate. (F) Percent neutralization of antigenically diverse HCVpp by HCABs and E1-dependent reference mAbs AR4A, IGH526, and HEPC112 at 100 µg/mL. VSV-Gpp is included as a negative control for non-specific neutralization. IC_50_ potency was measured using serial mAb dilutions and gt1a strain Bole1a HCVpp. mAbs that did not reach 50% neutralization at the highest mAb concentration tested (100 µg/mL) were assigned an IC_50_ of 200 µg/mL. Neutralization was measured in duplicate and represents an average of between 1 and 5 independent experiments. (G) E1E2 binding competition of 11 HCABs and 8 E1-independent (green) or E1-dependent (blue) reference mAbs. Binding of each biotinylated antibody to Bole1a E1E2 lysates was assessed in the presence and absence of each non-biotinylated blocking antibody. Darker purple indicates greater % competition. Black boxes indicate clusters. Data represents one experiment tested in duplicate for most combinations. For combinations tested in two or more experiments, results were averaged.

Isolation of E1-dependent mAbs was facilitated by the recent development of an engineered soluble E1E2 glycoprotein heterodimer with C-terminal synthetic zipper (SZ) domains, designated sE1E2.SZ^36,43–45^. We optimized a new protocol to stain B cells with purified, Avi-tagged^46^, biotinylated sE1E2.SZ (sE1E2.SZ-Avi-Bio) probes derived from strain H77, conjugated to two distinct fluorophores, as well as antibodies against B cell markers to identify class-switched B cells (CSBC: CD19 or 20+, IgM-,IgD-) (Fig. 1B). We then flow-sorted single E1E2-reactive CSBC from a C117 sample collected 45.3 months after acute infection, and 31 months after spontaneous HCV clearance. CSBC were sorted into individual wells in a 96 well plate and cultured for 14 days with irradiated 3T3-msCD40L cells, CpG, BAFF, IL-2, IL-21, and IL-17 to stimulate expansion, plasmablast differentiation, and antibody production^47^. The majority (67%) of wells produced detectable IgG in supernatants after stimulation, and we measured binding of these IgG+ supernatants to purified, soluble E2 (sE2) proteins and lysates of E1E2-transfected cells (Fig. 1C). Both sE2 and E1E2 lysates were derived either from strain 1a154 (H77) or a mixture of three other antigenically diverse HCV strains (1a157, 1b09, 1b21^11^). We found that 108 of 134 (81%) HCV-reactive single B cell culture supernatants bound sE2 (i.e., binding was E1-independent), and 25 of 134 (19%) of supernatants bound E1E2 lysates, but not sE2 (i.e., binding was E1-dependent). We obtained both immunoglobulin heavy chain (IgH) and light chain (IgL) variable sequences from 15 (60%) of these E1-dependent cultures. We successfully synthesized, cloned, and expressed 11 of these authentically paired IgH and IgL sequences to generate recombinant mAbs, which we designated hepatitis C antibodies (HCABs) (Fig. 1D). Analysis of the variable sequences of these mAbs demonstrated that each was derived from a distinct clonal lineage (Table S1). To confirm that the recombinantly expressed mAbs replicated the supernatant results, we tested binding of the mAbs to sE2 and E1E2 lysates, along with known E1-independent reference mAb HCV1^48^ and two E1-dependent reference mAbs, AR4A^34^ and IGH526^49^. As expected, all HCABs bound E1E2 and not sE2, with the exception of HCAB109, which did not bind well to either sE2 or E1E2 lysates (Fig. 1E). We next tested the binding affinity of these HCABs to the lysates of cells transfected with a neutralization sensitive, computationally-derived, ancestral genotype 1a E1E2 variant, Bole1a^50^. All HCABs had 50% effective binding concentrations (EC_50_s) of less than 1 µg/mL, with the exception of HCABs 105 and 109, which had EC_50_s greater than 10 µg/mL (Fig. S2). Due to the low binding affinity of HCABs 105 and 109, we excluded them from subsequent experiments.

We next tested the ability of these E1-dependent HCABs to neutralize the antigenically diverse HCVpp panel^23^ (Fig. 1F). As positive controls, we included reference E1-dependent neutralizing mAbs AR4A, IGH526, and HEPC112. As a negative control against non-specific neutralization, we included VSV-Gpp, a pseudoparticle made using the envelope protein of vesicular stomatitis virus^51^. We used the true-positive cutoff for neutralization (>25%), which was established in prior studies using this same HCVpp panel^23,27^. The majority of the 11 HCABs neutralized multiple HCVpp in the panel, and seven of them (64%) neutralized more than half of the panel, which is the threshold used to define bNAbs^27^. Reference mAbs AR4A, IGH526, and HEPC112 showed the expected phenotypes, with AR4A exhibiting very broad neutralization (14/15 HCVpp)^23^, IGH526 showing broad neutralization (12/15 HCVpp)^32^, and HEPC112 exhibiting narrow neutralization (4/15 HCVpp)^31^. Neutralization of VSV-Gpp by all mAbs was below the 25% positive cutoff. To determine the potency of neutralization, we measured the ability of serial dilutions of each mAb to neutralize HCVpp produced with Bole1a E1E2. The potency of the HCABs was variable, ranging from 0.4 to greater than 100 µg/mL, with the most potent HCABs demonstrating potency similar to AR4A.

To define the binding sites of these novel HCABs relative to previously described antigenic sites using binding competition experiments, we selected a panel of E1-independent and E1-dependent reference mAbs that bind all known sites across E2 and E1. HCV1 is an E1-independent bNAb that binds to AS412, a linear epitope near the N-terminus of E2^48^. HEPC74 is another E1-independent bNAb that binds to the front layer of E2^13,52^. HCAB7 and HCAB5 are also E1-independent mAbs that bind to the beta sandwich and back layer of E2, respectively^27^. AR4A is an E1-dependent bNAb that binds to the AR4 site in the bridging domain of E2^35,53^. HEPC111 is another E1-dependent mAb that binds to AR5, an E1-dependent site on E2 that is non-overlapping with AR4^31^. HEPC112 is a narrow-breadth neutralizing antibody that has been shown by alanine scanning mutagenesis to have critical binding residues in E1^31,54^. IGH526 is one of three highly related bNAbs (along with IGH505 and IGH520) that binds to a conserved alpha helix in the stem of E1^35,44,49^.

We used this comprehensive reference mAb panel in E1E2-binding competition assays with the 11 novel E1-dependent HCABs. In these assays, E1E2 was first incubated with a high concentration of a blocking mAb, followed by serial dilutions of a second, biotinylated mAb, which was then detected with streptavidin-horseradish peroxidase. For each pairwise combination of mAbs, each mAb was tested both as the blocking mAb and as the second, biotinylated mAb. Based on competition between each pair of antibodies, the eight reference mAbs and eleven novel HCABs clustered into four competition groups (Fig. 1G). As expected, the E1-independent mAbs HCV1, HEPC74, and HCAB7 did not compete with any E1-dependent HCABs. The first competition group included reference mAbs HCAB5 and HEPC111, and novel E1-dependent HCAB103. HCAB5 is an E1-independent bNAb that was previously shown to compete with HCAB40, a back layer mAb (PDB: 8W0X)^27^, and HEPC111 is an E1-dependent mAb that probably also binds to the E2 back layer, based on alanine scanning analysis^31^. We have also shown that HEPC111 competes with AR5A^34^, which is also a back layer binding, E1-dependent bNAb (Fig. S3). Together, these data indicate that HCAB103 likely binds at the back layer of E2 in an E1-dependent manner. The second and third competition groups partially overlapped, with E1-dependent reference mAb IGH526 exhibiting competition with some members of each group. The second competition group included IGH526, E1-dependent reference mAb AR4A, and novel E1-dependent HCABs 104 and 114. Notably, although AR4A and IGH526 both competed with HCABs in this group, they did not compete with each other, as expected, since they bind to non-overlapping epitopes in recent cryo-EM structures^35,36^. The third competition group included IGH526 and novel E1-dependent HCABs 110, 111, and 100. The fourth competition group included E1-dependent reference mAb HEPC112 and HCABs 102 and 115. No structure exists for HEPC112, but previous alanine scanning analysis showed critical binding residues in the N-terminal domain and putative fusion peptide (pFP)-containing region (PCR) of E1^35,54^, indicating that HCABs 102 and 115 likely bind to E1. Taken together, these data indicate that the 11 novel E1-dependent HCABs bind to at least four distinct antigenic sites, with these sites at least partially overlapping with previously described AR4, AR5, IGH526, and HEPC112 binding sites.

### AlphaFold 3 confidence in mAb-E1E2 structure predictions is significantly improved by the use of diverse E1E2 variants

We hypothesized that we could use AlphaFold 3 (AF3), a generative artificial intelligence (AI) that predicts the molecular structures of multi-protein complexes, to define the binding epitopes of E1-dependent HCABs at higher resolution. Given the genetic diversity of HCV E1E2^55–61^ and the importance of co-evolutionary relationships in the multiple sequence alignment (MSA) that AF3 embeds before inference^37^, we hypothesized that AF3 confidence would be improved through the generation of predictions using diverse E1E2 sequences. We compiled a panel of 151 genetically and antigenically diverse E1E2 variants from six HCV genotypes^23^, with each variant differing at an average of 23% of amino acids (Fig. 2A). To assess the impact of this E1E2 sequence diversity on AF3’s prediction confidence, we generated AF3 predictions of antibody variable fragment (Fv)-E1E2 complexes for 22 HCV mAbs, including reference mAbs and novel HCABs. We compared the maximum AF3 chain-pair interface predicted template modeling (iPTM) confidence score for the interface between the immunoglobulin heavy chain and either E1 or E2 (IgH-E1/E2 iPTM) of each Fv obtained with 1000 random seeds (5000 predictions) using only a single E1E2 variant (AMS0232^35^) or with 1000 seeds distributed across the diverse E1E2 panel. Maximum IgH-E1/E2 iPTM of each Fv was significantly greater with the diverse E1E2 panel than with the single E1E2 variant (Fig. 2B). We next ran additional independent random seeds for each Fv-E1E2, distributed across the diverse E1E2 panel, generating as many predictions as possible for each complex, given available computational time (median of 74,000 seeds per Fv). As expected, the maximum IgH-E1/E2 iPTM also improved significantly with a greater number of seeds (Fig. 2B). We assessed the maximum IgH-E1/E2 iPTM for each Fv-E1E2 variant combination using the 1000 seed per Fv-E1E2 dataset (Fig. 2C). iPTMs were ≥ 0.6 across the largest number of E1E2 variants for HEPC74^52^, AR3A^21^, HEPC3^13^, HCAB64^27^, and HCAB55^27^ (124-146 variants for these mAbs vs. 0-31 variants for the other reference mAbs). Notably, these five mAbs all bind to the E2 front layer. Antigen binding fragment (Fab)-E2 structures of three of these mAbs (HEPC74, AR3A, and HEPC3) were published before the AF3 training cutoff date, which may explain the high-confidence AF3 predictions for these structures, but structures of HCAB64 and HCAB55 were published after the AF3 training cutoff date. The increase in IgH-E1/E2 iPTM with the use of the diverse E1E2 panel was not driven by any single E1E2 variant, as maximum iPTM values for the set of mAbs did not differ significantly across variants (p=NS) (Fig. 2C). To assess the sensitivity of AF3 at different iPTM thresholds, we analyzed the number of mAbs with Fv-E1E2 predictions reaching iPTM thresholds ranging from 0 to 1, using the AMS0232 1000 seed, the diverse E1E2 1000 seed, and the diverse E1E2, ∼74,000 seed datasets (Fig. 2D). As expected, with all three datasets, the number of mAbs meeting the iPTM threshold dropped as the threshold increased. The largest number of mAbs met each iPTM threshold using the diverse E1E2, ∼74,000 seed dataset. Using this dataset, we specifically considered the number of mAbs with predictions meeting iPTM thresholds ≥ 0.6 or ≥ 0.8, since predictions meeting these thresholds had previously been shown to be accurate or highly accurate, respectively^37^. Of 22 mAbs, predictions for 21 reached the 0.6 iPTM threshold, but only 11 reached the 0.8 threshold. Together, these data show that AF3 confidence in Fv-E1E2 predicted structures is significantly improved by the use of diverse E1E2 variants, and by increasing the number of seeds. After maximizing iPTMs, most mAbs (21/22) reached an iPTM threshold of ≥ 0.6, but only half of the mAbs reached an iPTM threshold of ≥ 0.8, indicating that this higher iPTM threshold is probably too stringent for practical use with Fv-E1E2 structure predictions.

**Fig 2.**
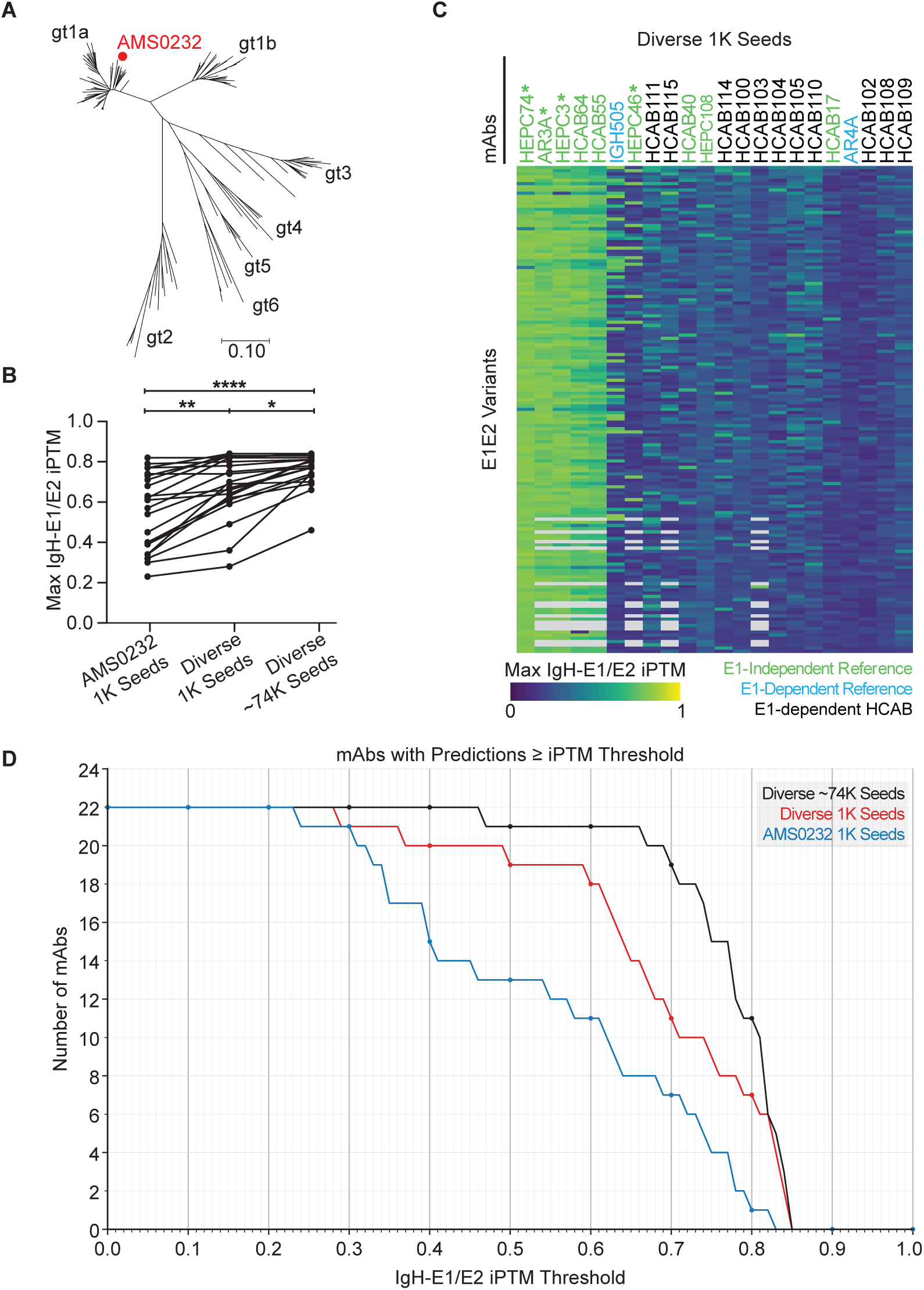
AlphaFold 3 prediction confidence is significantly improved by the use of diverse E1E2 variants, and the effect is not driven by any individual variant. (A) Maximum-likelihood phylogenetic tree of the 151 E1E2 amino acid sequences used for AF3 predictions. The tree was generated in MEGA 12 using the Jones-Taylor-Thornton model with gamma distribution and a proportion of invariant sites. Genetic distance is drawn to scale, and genotypes are labeled. The red dot indicates strain AMS0232 E1E2. (B) Maximum value of IgH-E1/E2 iPTM of Fv-E1E2 complexes of 22 reference mAbs and HCABs across three AF3 datasets, obtained using strain AMS0232 only or the diverse E1E2 panel, with the indicated number of seeds. Groups were compared using Friedman’s non-parametric test with Dunn’s correction for multiple comparisons. **** p<0.0001, ** p<0.01, * p<0.05. (C) Maximum IgH-E1/E2 iPTM by mAb-E1E2 variant combination for the 1000 seed per complex dataset using the diverse E1E2 variant panel. Colors represent the maximum value for that combination: yellow/green is higher iPTM, blue is lower iPTM. HCABs (black) and E1-independent (green) or E1-dependent (blue) reference mAbs are arranged from left to right from highest to lowest number of variants with a maximum iPTM > 0.6, and E1E2 variants are arranged from top to bottom from highest to lowest median value across all mAbs. Reference mAbs with experimental structures published before the AF3 training cutoff date are marked with an asterisk. (D) The number of mAbs with maximum IgH-E1/E2 iPTM values greater than or equal to each iPTM threshold from 0 to 1, using three AF3 datasets.

### iPTM ≥ 0.6 is a poor predictor of accuracy of AF3 predicted structures

AF3 predictions with high iPTM confidence scores yet definitively incorrect structures, termed hallucinations, are particularly difficult to detect and avoid for antibody-antigen complexes^38^, so we next tested the accuracy of AF3 predicted structures of HCV mAbs by comparing previously published, experimentally-determined Fab-E1E2 or Fab-E2 structures of 11 HCV mAbs^13,21,27,35^ to AF3 predictions of the Fvs of the same mAbs in complex with full-length E1E2 (66,120-1,326,455 AF3 predictions per Fv-E1E2 complex). Seven of the eleven experimental reference structures were published after the AF3 training cutoff date, so could not have been included in the training dataset. To assess the presence of hallucinations in our Fv-E1E2 predicted structures, we used DockQ, a standard metric for measuring the quality of protein-protein interface predictions^62,63^. The size and sequence differences of the AF3 predictions relative to their corresponding experimental structures necessitated modifications to the standard DockQ pipeline. Since all AF3 predictions contained Fvs and full-length E1E2, while some experimental structures were obtained with Fabs and/or E2 only, we first trimmed each pair of AF3 and experimental structures to only those residue positions that existed in both structures, prior to DockQ calculation. Based on prior studies showing bonding only with E2 for AR4A and only with E1 for IGH505, only the E2 chain was used for AR4A and only the E1 chain for IGH505 DockQ calculations. We also merged heavy and light chains together into one Fv chain for calculation to assess the complete Fv-E1E2 interface, regardless of whether binding of a particular Fv was more heavily weighted to either the heavy or light chain. We then plotted the relationship of this accuracy metric (Fv-E1/E2 DockQ) with the AF3 confidence score (IgH-E1/E2 iPTM) for each AF3 predicted structure of the eleven reference mAbs (Fig. 3).

**Fig 3.**
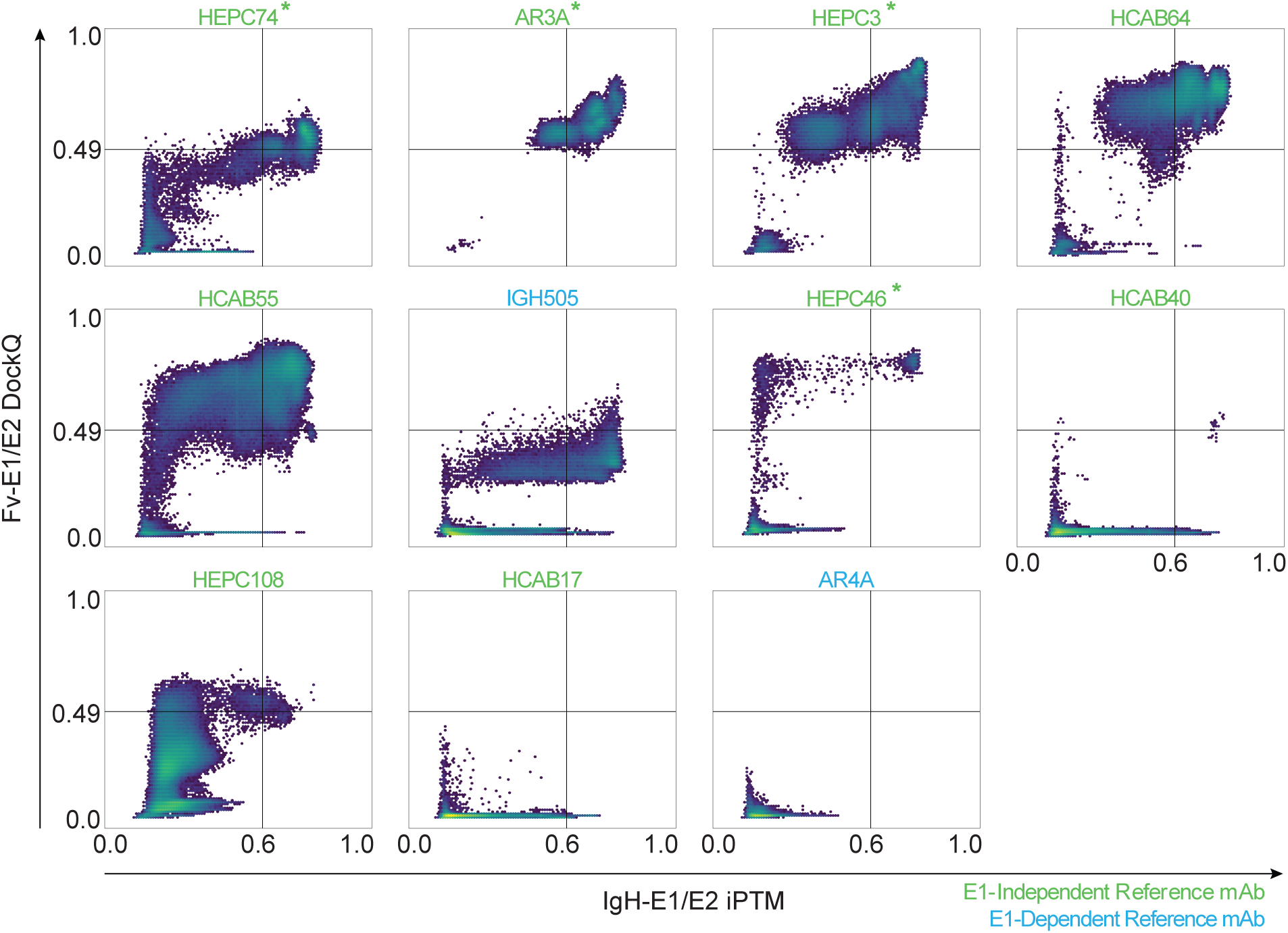
iPTM ≥ 0.6 is a poor predictor of accuracy of AF3 predicted structures. Hexbin heatmaps of all AF3 predictions for E1-independent (green) and E1-dependent (blue) HCV reference mAbs, correlating IgH-E1/E2 iPTM and Fv-E1/E2 DockQ score. Dividing lines indicate high AF3 confidence (iPTM ≥ 0.60) and high accuracy (DockQ ≥ 0.49). Color of each hexbin shows the number of predictions from blue (1 prediction) to yellow (10^5^ predictions). Reference mAbs with experimental structures published before the AF3 training cutoff date are marked with an asterisk.

In prior publications^37,62,63^, DockQ scores were divided into inaccurate (<0.23), acceptable (0.23-<0.49), medium (0.49-<0.8), and high quality (≥0.8). For this study, we used these same definitions for inaccurate and acceptable models but defined all models with DockQ scores ≥ 0.49 as highly accurate^62^. Since some predictions for most mAbs reached an iPTM threshold ≥ 0.6, which was predictive of reasonable accuracy in a prior study^37^, we focused our analysis on predictions with iPTM values above this threshold. None of the predictions for reference mAb AR4A exceeded an iPTM of 0.6. In contrast, mAbs HEPC74, AR3A, HEPC3, and HEPC46 showed many predictions with iPTM >0.6, with a strong correlation between iPTM and DockQ, potentially reflecting their possible inclusion in the AF3 training dataset. Above an iPTM cutoff of 0.6, all Fv-E1E2 predictions for HEPC74, AR3A, HEPC3, HEPC46, and HEPC108 were acceptable (DockQ > 0.23) or highly accurate (DockQ > 0.49). However, above this same iPTM cutoff, mAbs HCAB64, HCAB55, IGH505, and HCAB40 showed some hallucinations, with a mixture of highly accurate, acceptable, and inaccurate predictions. All predictions for mAb HCAB17 were inaccurate, including predictions with iPTM > 0.6. Overall, although predictions exceeding the 0.6 iPTM threshold for some mAbs were highly accurate, many predictions for some mAbs were inaccurate despite exceeding this iPTM threshold.

### 3D-spatial clustering along with iPTM identifies accurate AF3 predictions

Since iPTM ≥ 0.6 was insufficient to identify accurate AF3 predictions, we hypothesized that repeated high-iPTM predictions showing an Fv at the same E1E2 binding site would more accurately identify correct Fv-E1E2 predictions. To test this hypothesis, we developed a method to spatially map the Fv binding site in three dimensions relative to E1E2 for each AF3 predicted structure. For each AF3 prediction of Fv-E1E2, we aligned E1E2 against a recently published cryo-EM structure of E1E2^35^ in three different reference frames (E1, E2, and E1E2). For each prediction with IgH-E1/E2 iPTM ≥ 0.4, in each reference frame, we calculated the center of mass (COM) of the Fv, mapping a spatial location for Fv binding relative to E1E2 (Fig. 4). For each reference mAb, for a median of about 43,000 predictions, we mapped the spatial Fv locations relative to E1E2 and used DBSCAN^64^ to identify Fv clusters. The alignment reference frame (E1, E2, or E1E2) that produced the fewest clusters was chosen for further analysis.

**Fig 4.**
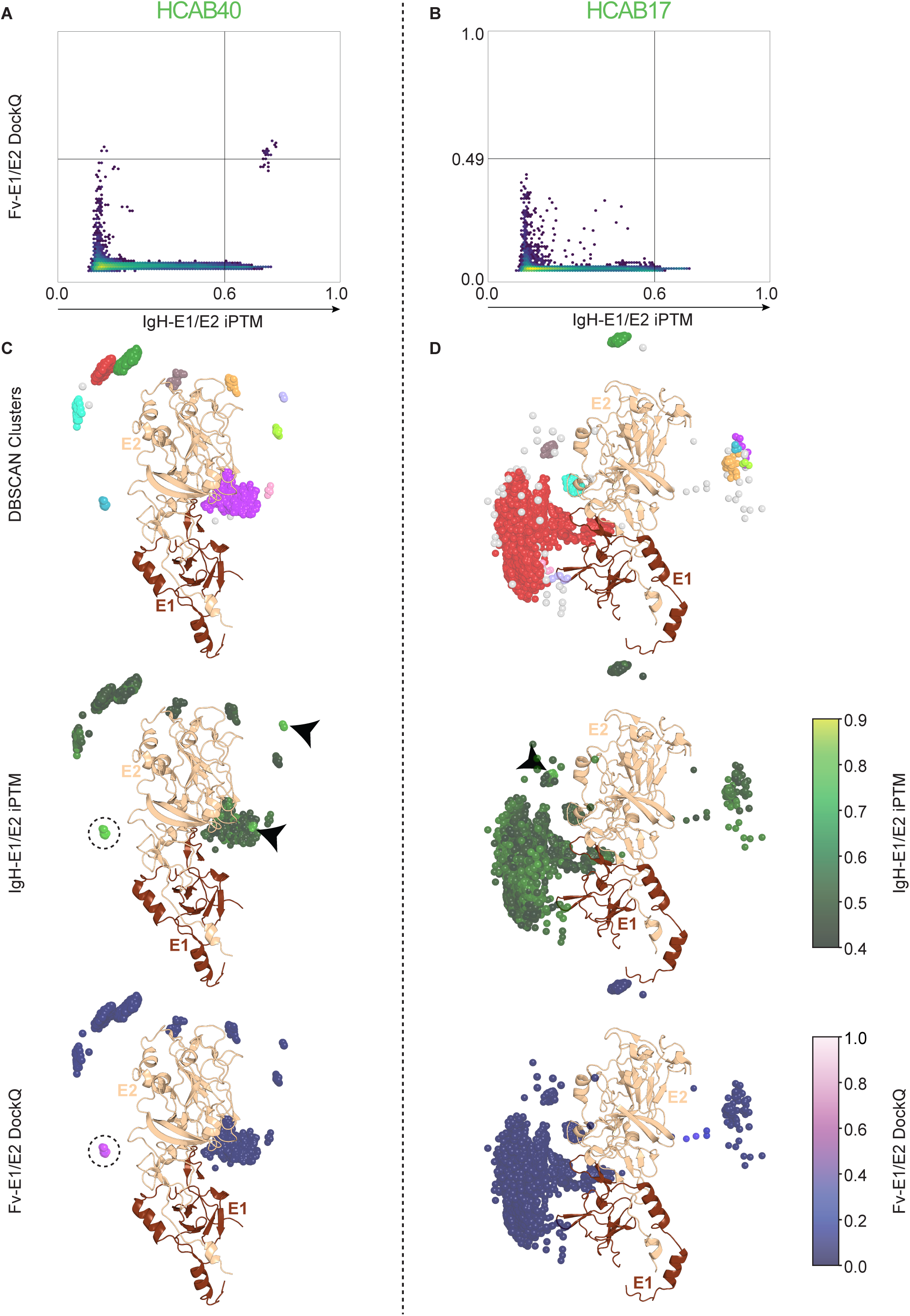
3D-spatial clustering of AF3 predictions with iPTM (mAbClust) allows identification of accurate epitope predictions. (A) Hexbin heatmap from Fig. 3 of iPTM vs. DockQ for reference mAb HCAB40. (B) Hexbin heatmap from Fig. 3 of iPTM vs. DockQ for reference mAb HCAB17. (C-D) Fv centers of mass (COMs) located in 3-dimensional space for all predictions with IgH-E1/E2 iPTM ≥ 0.4 of HCAB40-E1E2 (N=14,464) (C) or HCAB17-E1E2 (N=16,060) (D). Top panels show predictions colored categorically by clusters identified by DBSCAN. Predictions not belonging to any DBSCAN cluster are colored gray. Middle panels show the same predictions, with COM spheres colored by the IgH-E1/E2 iPTM from dark green (0.4) to light green (0.9). Circled cluster has the highest median iPTM for HCAB40 (0.77). Arrowheads indicate the location of high iPTM predictions in low median iPTM (<0.68) clusters, or high iPTM predictions in clusters with fewer than 11 predictions. Bottom panels show the same predictions, with COM spheres colored by their Fv-E1/E2 DockQ score from dark blue (low) to light purple (high). The same circled HCAB40 cluster has a median DockQ of 0.52 (highly accurate). All locations are shown in relation to the structure of E1E2 from PDB 7T6X^35^, with E2 colored in tan and E1 in brown.

For each reference Fv-E1E2, we calculated the median iPTM and median DockQ of each spatial cluster, observing that the median iPTM of all predictions in each spatial cluster correlated with the accuracy (DockQ) of the predictions in that cluster. To illustrate this, we compared HCAB40 and HCAB17, two antibodies with a large number of high iPTM, incorrect predictions (i.e., hallucinations). Despite many hallucinations, HCAB40 (Fig. 4A) did have some highly accurate (DockQ > 0.49) predictions, while all HCAB17 predictions (Fig. 4B) were inaccurate (DockQ < 0.23). In the spatial plot of HCAB40 predictions, 30 high-iPTM predictions clustered together, resulting in a cluster with a high median iPTM (0.77). Other high-iPTM predictions formed small clusters containing fewer than 11 predictions, or clustered with low-iPTM predictions, resulting in clusters with lower median iPTMs (Fig. 4C). The high median iPTM cluster also had a high median DockQ score (0.52), indicating that the predictions in the cluster were accurate. For HCAB17, all high-iPTM predictions clustered with low-iPTM predictions, resulting in no clusters with a high median iPTM (Fig. 4D). None of these clusters had high DockQ scores, indicating that none of the predictions were accurate.

To determine an optimal spatial cluster median iPTM cutoff to identify clusters containing accurate predictions, we examined the spatial cluster information for all reference mAbs. We first excluded clusters containing fewer than 11 predictions and then plotted the correlation between median iPTM of each cluster and the median DockQ value of that cluster (Fig. 5A). We chose a threshold that was below the median iPTM of all highly accurate clusters (median DockQ > 0.49) and above the median iPTM of all inaccurate clusters (median DockQ < 0.23) as the minimum median iPTM cutoff to identify accurate clusters (cutoff = 0.68). To choose a single predicted structure of each Fv-E1E2 for detailed analysis, we selected the cluster of predictions with the highest median IgH-E1/E2 iPTM above the cutoff and computed the spatial centroid of that cluster. We then calculated the distance of each prediction in the cluster to the cluster centroid. The prediction with the highest IgH-E1/E2 iPTM among the 10 closest predictions to the cluster centroid was chosen as the best predicted structure for each Fv-E1E2 complex. In summary, this algorithm for selection of accurate AF3 predictions, which we call mAbClust, includes spatial clustering of Fvs from the ensemble of predictions, identification of clusters with highest median iPTM values that exceed a minimum threshold, and selection of highest iPTM predictions near the centroid of these high median iPTM clusters.

**Fig. 5.**
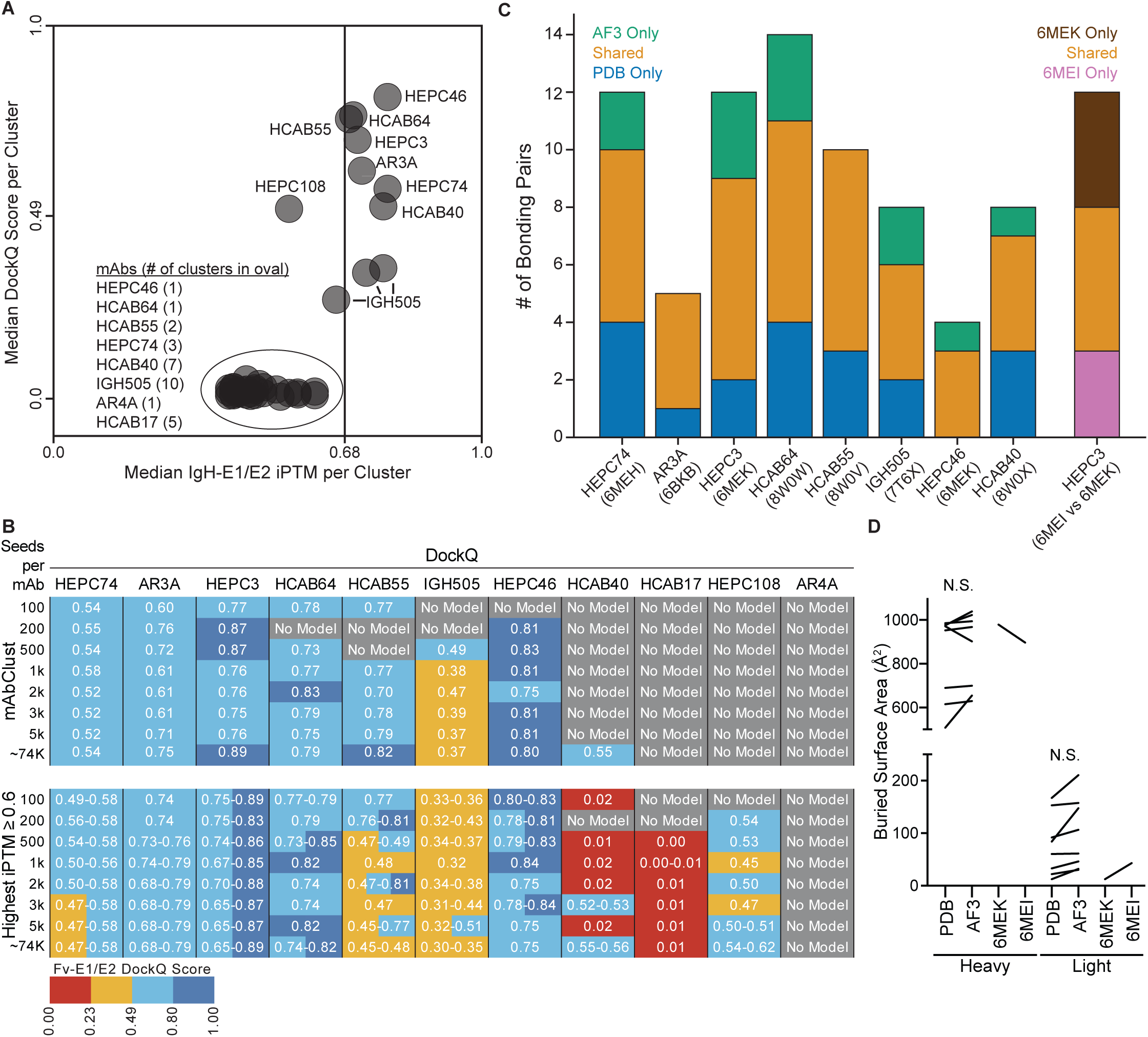
mAbClust selection of AF3-predicted Fv-E1E2 structures allows accurate mapping of atomic interactions. (A) Correlation of the median IgH-E1/E2 iPTM of each spatial cluster with the median Fv-E1/E2 DockQ of the same cluster for 11 reference mAbs. Vertical line indicates the median iPTM threshold selected to identify clusters containing accurate predictions. (B) HBPlus hydrogen-bonding analysis of Fv-E1E2 residue pairs forming hydrogen bonds in mAbClust-selected AF3 predictions and corresponding experimental structures (PDB). Bonds were considered to be shared if the positions of bonding residue pairs matched between AF3 and experimental structures. The bar on the far right shows the same analysis using two experimental X-ray crystal structures of HEPC3-E2 complexes obtained with different E2 variants. (C) DockQ scores of predictions selected either using mAbClust (top table) or by selecting the prediction(s) with the highest iPTM ≥ 0.6 (bottom table). Prediction datasets were generated using the diverse E1E2 panel, ranging in size from 100 to ∼74,000 predictions. Red indicates incorrect predictions (DockQ < 0.23), orange indicates acceptable predictions (0.23 ≤ DockQ < 0.49), and blue indicates highly accurate predictions (0.49 ≤ light blue < 0.80 ≤ dark blue). (D) Buried surface area (BSA) of heavy and light chains of each Fv in complex with E1E2, comparing mAbClust-selected AF3 predictions to the corresponding experimental structures (PDB). Statistical comparisons use repeated measures one-way ANOVA with Sidak’s correction for multiple comparisons (p=NS, not significant). Comparisons between the two X-ray crystal structures of HEPC3 (6MEK and 6MEI) are also shown.

To evaluate the overall performance of the mAbClust algorithm using datasets ranging from 100 seeds per mAb to a median of ∼74,000 seeds per mAb, we calculated the Fv-E1/E2 DockQ scores for AF3 predictions of each reference mAb selected either by the standard approach based only on the highest iPTM score ≥ 0.6, or using the mAbClust algorithm (Fig. 5B). Both approaches correctly determined that no accurate predictions were available for mAb AR4A. Notably, for HCAB17, use of highest iPTM ≥ 0.6 selected inaccurate predictions from all datasets of 500 seeds or greater. Similarly, for HCAB40, use of highest iPTM ≥ 0.6 selected inaccurate predictions from the 100, 500, 1000, 2000, and 5000 seed datasets. In contrast, the mAbClust algorithm avoided these inaccurate predictions, correctly determining that no accurate predictions were available in any of the HCAB17 datasets or in the HCAB40 100, 500, 1000, 2000, 3000, and 5000 seed datasets, while correctly identifying a highly accurate HCAB40 prediction in the ∼74,000 seed dataset. For HEPC108, use of highest iPTM ≥ 0.6 correctly selected accurate predictions from datasets of 200, 500, 2000, 5000 and ∼74,000 seeds, and acceptable predictions from the 1000 and 3000 seed datasets, while mAbClust incorrectly determined that no accurate predictions were available. For the remaining seven reference mAbs, both highest IPTM ≥ 0.6 and mAbClust methods selected accurate predictions, with DockQ scores generally higher for the mAbClust-selected predictions. Overall, mAbClust improved DockQ scores for most mAbs, while demonstrating a critical advantage over highest iPTM ≥ 0.6 in rejecting all inaccurate, hallucinated AF3 predictions.

To determine whether these predictions could be used to describe antibody:antigen interfaces at atomic resolution, we compared the predicted hydrogen bonds between Fv and E1E2 in mAbClust-selected AF3 predictions to the bonds in experimental structures of the same mAbs in complex with E2 or E1E2 (Fig. 5C, Tables S2 to S9). All corresponding AF3 and experimental structures shared Fv-E1E2 bonding residue pairs, ranging from 50% of all pairs (HCAB40, HCAB64, HEPC74, and IGH505) to 80% of all pairs (AR3A). We suspected that some variation between AF3 predictions and experimental structures could be due to the use of different E1E2 variants in AF3 and experimental structures. To assess how much bonding variation would be expected between experimental structures that also used different HCV variants, we completed this same analysis with two separately deposited high resolution crystal structures of the same antibody (HEPC3) in complex with either strain 1a53 or 1b09 E2 (Fig. 5C and Table S10). Less than 50% of Fv-E2 paired bonding residues were shared between the two experimental structures. This analysis confirmed that some variation in bonding patterns between AF3 predictions and experimental structures is expected and does not necessarily indicate that AF3 predictions are inaccurate. Aligning AF3 and experimental structures on E2 in PyMOL showed that AF3 correctly predicted not only the general location, but also the angle of approach and the placement and conformation of the Fv complementarity determining region (CDR) loops (Fig. S4), with root mean square deviation (RMSD) between CDRH3 of AF3-predicted and experimental structures ranging from 0.46-2.19 Å (Table S11). When comparing the CDRs of two crystal structures of HEPC3, the CDRH3 RMSD was 1.05 Å. The predicted Fv-E1E2 interfaces were also quantitatively similar, with no significant difference in buried surface area between each AF3 prediction and its corresponding experimental structure (p=NS) (Fig. 5D). Taken together, these data indicate that mAbClust identifies atomically correct AF3 predictions of HCV Fv-E1E2 complex structures more accurately than highest iPTM score ≥ 0.6, while rejecting inaccurate, hallucinated predictions.

### E1-dependent, broadly neutralizing HCAB 104 binds to a novel quaternary epitope, with bonds to both E1 and E2

Next, we used AF3 and mAbClust to generate and select a high confidence prediction for the structure of the Fv of HCAB104 in complex with E1E2. We selected this HCAB for further analysis because of its unexpected binding competition with both AR4A and IGH526. We generated 1,272,970 HCAB104 Fv-E1E2 predictions and observed spatial clusters as we had previously with reference mAbs (Fig. 6A). One of these clusters (N=45 predictions) had a median iPTM of 0.75, indicating that the predictions in that cluster were likely to be accurate (Fig. 6B). We selected the highest iPTM prediction near the centroid of this high median iPTM cluster for subsequent analyses. The density of the HCAB104 Fv was evenly split between E1 and E2 (Fig. 6C). When the experimentally determined Fv structures of AR4A and IgH505 (PDB 7T6X) were merged with the HCAB104-E1E2 predicted structure by aligning on E2 (for AR4A) and E1 (for IGH505), we could see a progression of density from AR4A to HCAB104 to IGH505, with clashes between AR4A and HCAB104 and between HCAB104 and IGH505 (Fig. 6D). This was consistent with the observed binding competition of both AR4A and IGH526 with HCAB104, and with no observed competition between AR4A and IGH526. Remarkably, the CDRH3 loops of HCAB104 and AR4A overlapped (Fig. 6E), consistent with a recent report describing structural homology of E2 bridging domain mAbs^65^. The HCAB104 heavy chain made contacts with both E2 and E1, burying 681 Å^2^ and 317 Å^2^, respectively, while the light chain only contacted E2, burying 134 Å^2^ (Fig. 6F). The HCAB104 CDRH3 formed a complex network of interactions with the base of E2 that was highly similar to AR4A CDRH3 interactions with the same site (Fig. 6G and Table S13)^35,65^. This included hydrogen bonds to four of the same E2 residues (649, 695, 697, and 698) that are seen in the AR4A experimental structure. In addition to CDRH3 contacts, CDRH1 of HCAB104 formed two bonds with the stem of E2 (H33 – 701) and two with the C terminal end of E1 (H28 – 313, H30 – 318). These E1 contacts overlapped with the epitope of IGH505 (Fig. 6H and Tables S12-13). The light chain only formed a bond with the base of E2 (L91 – 648). The predicted binding residues for HCAB104 in E1 and E2 are highly conserved, with all but 1 residue more than 99% conserved across all HCV genotypes in the HCV-GLUE database (Fig. 6I). Taken together, these data indicate that HCAB104 represents a novel class of bridging domain bNAb targeting a highly conserved quaternary epitope including residues in both E1 and E2.

**Fig. 6.**
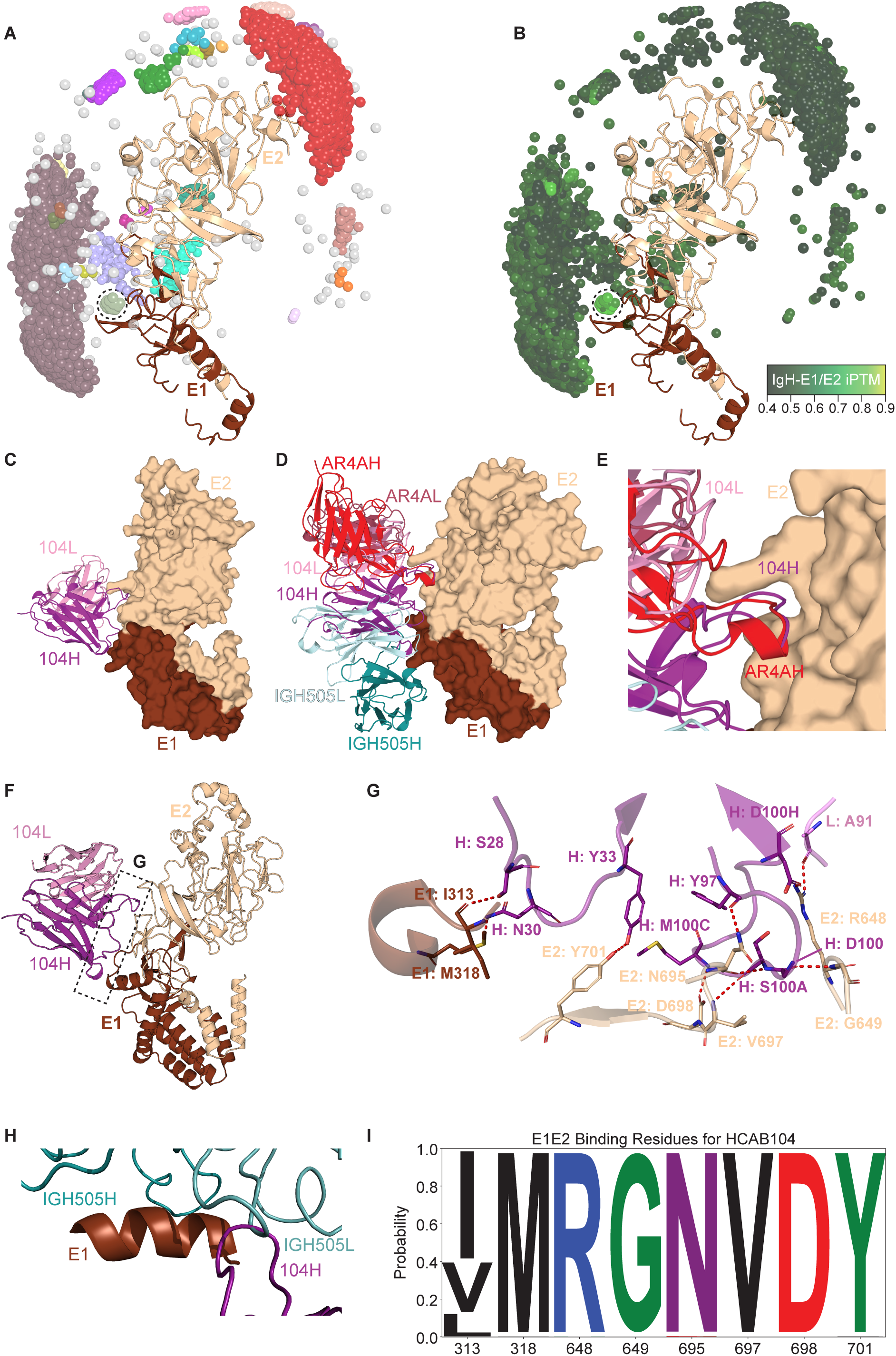
HCAB 104 binds to a novel quaternary epitope with bonds to both E1 and E2. (A) Fv centers of mass (COMs) in 3-dimensional space for all HCAB104 predictions with IgH-E1/E2 iPTM ≥ 0.4 (N=24,643) with 28 clusters identified by DBScan and 145 predictions not clustered (gray). (B) Fv COMs from A, colored by the IgH-E1/E2 iPTM from dark green (0.4) to light green (0.9). The circled cluster contains 45 predictions and has a median IgH-E1/E2 iPTM of 0.75. (C) The AF3 predicted, mAbClust selected Fv-E1E2 structure of HCAB104, with a ribbon diagram of the Fv (purple/pink) and a surface representation of E1E2 (E2 in tan, E1 in brown), showing binding at the interface between E1 and E2. 104H indicates the IgH, and 104L indicates the IgL. (D) Merging of the predicted structure of HCAB104 Fv-E1E2 with the experimental structure of AR4A (red) and IgH505 (cyan) in complex with E1E2 (PDB 7T6X), with alignment of AR4A on E2 and alignment of IGH505 on E1. (E) Detailed view of CDRH3 loops of HCAB 104 and AR4A in the E2-aligned structures, with HCAB 104 in purple/pink and AR4A in red. (F) Ribbon diagram of the HCAB104 Fv-E1E2 complex, with the region of interaction between Fv and E1E2 shown in a dashed box. (G) Detailed view of the dashed box in (F), showing HCAB 104 IgH and IgL bonds with E1 (brown) and E2 (tan). Antibody numbering uses the Kabat scheme, and E1E2 numbering is relative to strain H77. (H) Detailed view of the CDR loops of IgH505 (cyan) and HCAB 104 IgH (purple) interacting with E1 residues 314-324. IGH505 (PDB 7T6X) is aligned by E1 residues 314-324 to the same residues of the HCAB104 predicted structure. (I) Conservation of E1E2 residues predicted to form hydrogen bonds with HCAB 104. The height of each letter is scaled to the global frequence of that amino acid at that position in all sequences from all HCV genotypes in the HCV-GLUE database. E1E2 numbering is relative to strain H77.

## Discussion

There is a large unmet need for an effective HCV vaccine and compelling evidence that vaccine development is possible given the natural history of EN with clearance of genetically and antigenically diverse HCV infections. Mapping the epitopes targeted by EN with HCV clearance is essential for vaccine design. Although analyses of polyclonal plasma antibodies have indicated that E1-dependent bNAbs are associated with spontaneous control of HCV^2,18,42^, very few E1-dependent human mAbs have been isolated, and all were isolated from individuals with chronic HCV infection.

In this study, we developed a new protocol utilizing sE1E2.SZ to isolate E1E2-reactive B cells from an EN with spontaneous HCV clearance and carried out parallel supernatant screening and next generation single cell B cell receptor sequencing on the same B cells. We showed that binding of 19% of IgG+, HCV-reactive supernatants was E1-dependent. We expressed eleven E1-dependent mAbs and found that seven were bNAbs. Through binding competition ELISAs, we showed that the E1-dependent mAbs formed four distinct competition groups. We then developed and validated mAbClust, a new algorithm that improves identification of highly accurate AF3 predictions of antigen-antibody complexes. We used AF3 and mAbClust to generate a high-confidence predicted structure of a novel E1-dependent bNAb in complex with E1E2, showing that it binds to a quaternary epitope spanning E1 and E2. In agreement with binding competition data, this epitope is distinct from the epitopes of E1-dependent bNAbs isolated previously from persons with chronic HCV infection, which bind only to E2 or E1^21^. Together, these observations identify four E1E2 sites targeted in an EN with spontaneous HCV clearance, defining the structure of a quaternary bNAb epitope at one of those sites.

Current efforts to quality control or stabilize E1E2 vaccine antigens are often dependent on measurement of AR4A binding, since it binds to a conformational epitope at the E1E2 interface. These novel HCABs provide new tools to measure the conformational integrity of vaccine antigens at additional, biologically relevant bNAb epitopes. In addition, the near-identical binding mode of the CDRH3 loops of HCAB104 and AR4A, and the overlap of the HCAB104 epitope with the binding epitope of IGH505/520/526, identifies specific residues in E2 and E1 that are important bNAb targets. Since HCAB104 is predicted to bind to a quaternary epitope with contacts in both E1 and E2, we speculate that the mechanism of neutralization could involve interference with a conformational change at the E1E2 interface that is necessary for viral entry. Further work is needed to understand how bNAb binding at this interface site might affect conformational changes at the E1E2 interface or viral fusion with the host cell membrane.

Limitations of this study include the isolation of mAbs from a single individual, although it is an individual with highly desirable characteristics. In addition, a relatively small number of E1-dependent mAbs were isolated. Future studies should include application of this approach to isolate a larger number of mAbs from multiple participants with either spontaneous clearance or chronic infection to allow better comparison of mAb features associated with either outcome. An additional limitation is the reliance on a computationally predicted structure, rather than an experimental structure for high-resolution epitope mapping. We extensively validated the accuracy of the combined AF3 and mAbClust method for known HCV mAb-E1E2 complexes and saw that AF3 predictions were consistent with experimental structures and binding competition results. However, experimental structures remain the gold standard for structural epitope mapping. Future work will include additional validation of AF3-predicted structures using mutagenesis of putative binding residues and cryo-EM.

In this study, we examined the E1-dependent antibody repertoire in an EN who spontaneously cleared HCV without treatment, showing that E1-dependent bNAbs targeted four distinct sites on the E1E2 glycoprotein. We also used AF3 with mAbClust, a new algorithm that we developed, to generate a high-confidence predicted structure of a novel E1-dependent bNAb in complex with E1E2, showing that it binds to a quaternary epitope spanning E1 and E2. This novel bNAb epitope is distinct from the epitopes of E1-dependent bNAbs isolated previously from persons with chronic HCV infection. Together, these studies identify new neutralizing sites and a new bNAb epitope, advancing our understanding of mechanisms of HCV control and guiding HCV vaccine design. The AF3 with mAbClust approach for epitope mapping could have broad applications for other HCV mAbs and mAbs against other viruses.

## Materials and Methods

### Human participants

Baltimore Before and After Acute Study of Hepatitis (BBAASH) is a prospective cohort ^5,28,45–47^ of people who inject drugs in Baltimore who are followed from before the time they are infected with HCV, through spontaneous clearance and/or persistence of HCV. Participants in the BBAASH cohort are adults enrolled in Baltimore, MD^28^. These identity-unlinked samples are stored under code in aliquots in −80 freezers (plasma) or liquid nitrogen (PBMC) with continuous temperature monitoring. The study was approved by the Institutional Review Board of Johns Hopkins Hospital and informed written consent was obtained from all study participants.

### Cell lines

The Huh7 hepatoma cell line (male) was obtained from the JCRB cell bank. HEK-293T-CD81 knockout cells (female) were obtained from Dr. Joe Grove, University of Glasgow, United Kingdom. Expi293F (female) was obtained from ThermoFisher. 3T3-msCD40L cells (male) were obtained from the NIH AIDS reagent bank. Cells were grown under standard aseptic conditions at 37°C and authenticated based on expected microscopic morphology and phenotype (e.g. expected infection with HCVpp (Huh7) or high transfection efficiency and protein production (HEK-293T-CD81 knockout cells and Expi293F).

### Source of reference monoclonal antibodies

HEPC74, HCAB 5, and HCAB 7 were synthesized in the Bailey Laboratory as previously published^13,27^. HEPC111 and HEPC112 were provided by the James Crowe laboratory (Vanderbilt University)^31^. AR5A was purchased from Sigma Aldrich. AR4A was synthesized using the amino acid sequences published in PDB 7T6X. IGH526 was synthesized using the amino acid sequence published in PDB 4N0Y. HCV1 was synthesized using the amino acid sequence published in 4DGY.

### Soluble E2 variants used for B cell staining

Genes encoding 1a157, 1b09, and 1b21 E2 ectodomains, residues 384-643 (numbering based on strain H77), were cloned into a mammalian expression vector pHCMV3 that includes an N-terminal Ig-kappa secretion signal and expressed by transient transfection of Expi293F cells. Proteins were purified over 5 mL HisTrap HP columns on an Akta Pure 25L FPLC, concentrated, and polished using a Superdex 200 Increase gel filtration column, before desalting, buffer exchange, concentration, and site-specific biotinylation with BirA ligase (Avidity).

### E1E2-reactive B cell isolation

sE1E2.SZ-Avi-Bio probes were produced in the Thomas Fuerst laboratory (University of Maryland) as previously described^43^. Cryopreserved PBMC’s were thawed dropwise into warm RPMI-1640 containing 50% HI-FBS and 1:500 Benzonase, centrifuged at 400G for 5 minutes, counted, and rested overnight at 1-3E6/ml in 5% CO_2_ and 37 °C. Staining steps were carried out in PBS with 1% BSA and 2 mM EDTA, with washes in between. First, human anti-CD81 and Fc block were added to reduce non-specific binding to sE1E2 and flow antibodies respectively. Second, purified sE1E2-His-Avi-Biotin-Fluor was incubated with PBMCs at 2 µg per well. Finally, fluorescently labeled antibodies against CD3 and CD14 (B cell exclusion gate), CD19 and CD20 (B cell selection), IgM and IgD (to exclude unswitched B cells), and Propidium Iodide (viability) were added. Each of the above steps was carried out for 20-30 minutes in the dark. Cells were washed twice, passed through a 0.45 µm filter, resuspended and diluted, and run on a MoFlo flow sorter (BD).

### B cell stimulation and culture

Individual sorted B cells were stimulated and grown in culture with modifications to a standard protocol^47^. Briefly, 3T3-msCD40L cells were grown to confluence in T175 cell culture flasks before gamma irradiation and cryopreservation. On the day of the sort, culture media was prepared with IMDM + Glutamax, 10% heat-inactivated FBS, 1E5 irradiated cells/ml, 5µg/ml CpG, 0.1 µg/mL BAFF, 50 ng/ml IL-2, 50 ng/mL IL-21, and 1 ng/ml IL-17. The inner 60 wells of a 96-well plate were filled with 150 µl of this culture media and prepared for single cell sorting. After collection, plates were stored at 37 °C and 5% CO_2_ for 14 days. On day 14, plates were centrifuged at 500 g for 5 minutes, 100 µl of supernatant was moved to a V-bottom plate and frozen for analysis, and a cold mixture of 132 ml DEPC-H20, 1.7 ml RNase inhibitor, and 2 ml of 1 M Tris-HCl (pH 8.0) was prepared, 50 µl added to each well of cell pellet, briefly mixed by pipetting, before flash freezing and storage at −80 °C.

### Supernatant screening for IgG expression, sE2 and E1E2 specificity

Supernatant screening was performed in 384-well plates on unprocessed B cell culture supernatants as previously described^27^. Briefly, for measurement of intact IgG, anti-Fab antibody I5260 was adsorbed onto 384-well MaxiSorp plates for a sandwich ELISA with supernatant and detection by HRP anti-IgG (G18-145). For measurement of binding to sE2 or E1E2 cell lysates, GNA lectin was adsorbed onto the plate, followed by incubation with purified protein (5 µg/ml) or lysate (1:10 dilution), followed by supernatant, and detection with HRP anti-IgG (G18-145).

### B cell receptor sequencing

The lysed monoclonal cell pellets were thawed, and 4 µl of 1:400 diluted RNA was reverse transcribed into cDNA using random primers and SuperScript IV per the manufacturer’s instructions. For amplification of the variable regions of IgH, IgK, and IgL, 2.5 µl of cDNA was used in a 20 µl Kapa2G Fast Multiplex PCR with previously described nested primers^66^. The second round PCR primers were modified to include Ad 1.2 overhangs on the forward primers and Ad_min overhangs on the reverse primers. A third round of PCR was performed for 8-10 cycles, again using Kapa2G, in which unique molecular barcodes were added, as well as the Illumina P5 and P7 adapter sequences. Illumina MiSeq was used for next generation sequencing at 2×300 nt paired end reads. Sequences were aligned and annotated using MiXCR. Any wells whose most abundant sequence had fewer than 5 counts were discarded, and the top clone for each well was chosen for further analysis.

### Monoclonal antibody expression

Processed sequences were trimmed to the variable region, and any missing FR1 or FR4 amino acids were imputed from the germline. These were synthesized by Twist Biosciences and cloned into human IgG1 (pTwist CMV BetaGlobin WPRE Neo_IgG1Fc), IgK (pTwist CMV BetaGlobin WPRE Neo_Kappa_Tag), or IgL (pTwist CMV BetaGlobin WPRE Neo_Lambda_Tag) expression plasmids. After growth in NEB Stable E. coli cells, plasmids were purified using Zymogen Midi- or Maxiprep kits. Purified plasmids underwent long-read sequencing by Plasmidsaurus to prevent loss of insert, plasmid dimerization, or point mutations. Paired heavy and light chain plasmids were transfected into Expi293F cells at a 1:2 ratio. After 5-7 days, IgG was purified using HiTrap MabSelect PrismA 5 mL columns on an Akta Pure 25L or Masterflex Ismatec Peristaltic pump. Purified IgG was then desalted, concentrated, and buffer exchanged into PBS using 30K MWCO protein concentrators (Pierce). Purified IgG was sterile filtered before neutralization experiments.

### HCV envelope ELISAs and binding competition experiments

Binding competition experiments were carried out with each pair of mAbs, with each mAb tested as both the first, blocking antibody and the second, biotinylated antibody. Purified monoclonal antibodies were biotinylated using Thermo’s EZ-Link Micro NHS-PEG4 Biotinylation Kit and desalted. Immulon 2HB 96-well plates were coated with GNA lectin followed by saturating dilutions of Bole1a E1E2 cell lysates expressed in Expi 293F cells. After blocking in PBS + 1% goat serum+ 1% milk + 0.5% Tween-20, 20 µg/ml blocking antibody was added to appropriate wells. Each run of experiments included a non-blocked control for each biotinylated detection antibody to account for day-to-day variation. Biotinylated detection antibody was serially diluted five-fold from 20 µg/ml to 0.256 ng/ml. A second version of the protocol was developed during this work in which the second antibody was added after the blocking antibody without an intervening wash. This protocol increased the sensitivity of the assay and was performed for some repeat experiments. Detection was performed with HRP anti-Fc (IgG). All assays had at least two technical replicates. Biological replicates (different mAb production lots) when performed were averaged. Data were plotted in GraphPad Prism 10, area under the curve (AUC) was calculated, and % competition was defined as 1-(AUC with blocking antibody/AUC without blocking antibody). Background was defined as the average competition seen between pairs of reference antibodies with known non-overlapping experimental structures. Standard deviations above this background were used to bin percent competition values.

Binding measurements of supernatants or monoclonal antibodies to purified sE2 or lysates of cells transfected with E1E2 were carried out using the same protocol, omitting the blocking step and using HRP Anti-IgG for detection rather than Strepatividin-HRP.

### HCVpp expression and neutralization assays

HIV group-specific antigen (Gag)-packaged HCVpp were produced by lipofectamine-mediated transfection of HCV E1E2 expression plasmids, pNL4-3.Luc.R-E-expression plasmid, and pAdVantage (Promega) as previously described^23,27,67^. The panel of 15 HCVpp from Salas et al.^23^ was used in neutralization assays, as well as VSV-Gpp HCVpp^50^. Neutralization assays were performed as previously described^67^. The cutoff for true-positive neutralization was set at >25%. mAbs that did not reach 50% neutralization at the highest mAb concentration tested (100 µg/mL) were assigned an IC_50_ of 200 µg/mL.

### Source of E1E2 amino acid sequences for AlphaFold 3 predictions and sequence analysis

E1E2 sequences from Salas et al.^23^, Frumento et al.^7^ and additional sequences selected to represent genotypes 2-6 were downloaded from NCBI GenBank, LANL HCV^68^, and euHCVdb^69^ databases. A phylogenetic tree was generated in Mega 12. A sequence logo was generated in Python using the Logomaker package.

### Predicting Fv-E1E2 structures with AlphaFold 3

AlphaFold 3 weights were requested and granted from Google DeepMind. The databases and code repository were cloned onto two university-wide High Performance Computing clusters – the Rockfish cluster administered by ARCH, and the Discovery cluster administered by RIT. Inputs and outputs were generated and analyzed identically for both systems. A100, H100, and H200 GPU’s were utilized for prediction. Briefly, a database of HCV E1E2 sequences and Fab IgH and IgK/L variable domain sequences were maintained and programmatically assembled into input JSON’s with randomly chosen seeds. AF3 outputs – the model.cif file and summary_confidences.json were then parsed for statistical and structural analysis. Unless otherwise noted, all analysis was performed in Python.

### AF3 Processing

All AF3 models were aligned using the Pymol API to PDB 7T6X by E1, E2, and E1+E2. The center of mass of all atoms in the Fv was calculated in each of these three reference frames. Unless otherwise specified, anywhere we use iPTM, we are referring to the highest antigen heavy chain pair iPTM, which we define as the maximum value for each prediction between the two values: IgH-E1 iPTM (i.e. C/A chain-pair iPTM) and IgH-E2 iPTM (i.e. C/B chain-pair iPTM).

### DockQ Scoring

All models, both AF3 and reference PDB, were trimmed for DockQ analysis. Briefly, BioPython’s BLOSUM62 pairwise aligner was used to align the sequence of each chain of a given AF3 model with its counterparts of the reference PDB structure. The residue positions after alignment that did not have existing residues in either structure (i.e. unresolved or missing portions of the PDB structure, or constant regions that exist in the PDB structure but were not included in AF3 predictions) were removed. DockQ v2^62^ was installed from the Wallner Lab GitHub repository and run on all interfaces to extract the combined DockQ, the Heavy-Ag, Light-Ag, and Heavy-Light DockQ’s. In memory, Biopython’s PDB module was used to combine all heavy and light residues into a single chain, for Fv-E1/E2 DockQ analysis, following the approach described in their GitHub page.

### mAbClust Analysis

For each Fv-E1E2 complex, all predictions with iPTM ≥ 0.4 were filtered for clustering and the Fv XYZ COM coordinates were clustered by DBSCAN using scikit-learn. The alignment method (E1, E2, or E1E2) that produced the fewest clusters on the same data was carried forward. Clusters with fewer than 11 models were removed from the dataset, as were those with a median iPTM < the confidence cutoff. The top ranked cluster by median iPTM was chosen, the centroid of the cluster was calculated by mean of all coordinates in the cluster and the Euclidean distance of each model to this centroid was calculated using NumPy. The highest iPTM model amongst the 10 models in the cluster closest to the centroid was considered the chosen confident model of that Fv-E1E2 complex and used for further structural analysis.

### Computational Structure Analysis

Experimental and AF3-predicted structures were visualized and analyzed in Pymol (v3.1.5.1). Alignment of structures was performed using the cmd.align function. For CDR-RMSD analysis, AbNumber was used to annotate the IgG sequences. Each chain was superimposed against its corresponding experimental structure by residues in the framework region, after which the RMSD between the AF3 model’s CDRs and the corresponding CDRs of the PDB model was calculated. For bonding analysis, HBPlus v3.2 was used to predict hydrogen bonds with default parameters^70^. For buried surface area of interfacing regions, PISA v2.2.0 was used from the CCP4 software suite following their documentation^71^. GNU parallel was used for command-line high-throughput processing^72^. OpenMPI (v4.1.6) was used for multi-node parallel processing. Agentic coding agents were used for Python scripting and debugging. All AI output was reviewed manually.

## Acknowledgments

We thank the participants, staff, and faculty central to the BBAASH cohort as well as members of the Johns Hopkins Viral Immunity and Pathogenesis Center. We thank the Single Cell and Transcriptomics Facility of Johns Hopkins Medicine, as well as the Advanced Research Computing at Hopkins Facility and the Research IT@JH group, including Discovery HPC and SAFESTOR. The authors also wish to thank Nicole Frumento and Clinton Ogega for valuable discussions. Funding was provided by the National Institutes for Allergy and Infectious Diseases (NIAID) of the National Institutes for Health (NIH) R01AI127469 (to J.R.B. and A.I.F.) and U19AI159822 (to A.L.C., J.R.B., and A.I.F.).

## Author Contributions

H.T.P. and J.R.B. conceived the study. H.T.P, J.M, H.A., L.S.B., and H.Z. performed experiments. L.K., X.W., E.A.T, A.I.F, H.B.L., T.R.F, S.C.R, A.L.C., and J.R.B. contributed resources. K.K., H.Z., N.S., N.S., A.I.F., G.O., H.B.L., T.R.F, S.C.R, and J.R.B. contributed to methodology. H.T.P. and N.S. wrote software and performed computational analyses. H.T.P. and J.R.B. analyzed the data and wrote the first draft of the manuscript. H.T.P., J.M., H.A., L.S.B., L.K., X.W., K.K., H.Z., E.A.T., N.S., A.I.F., G.O., H.B.L., T.R.F., S.C.R., A.L.C., and J.R.B. interpreted the results and contributed to revisions of the final manuscript. J.R.B. supervised the project.

## Resource Availability

Code and molecular coordinates will be deposited in relevant repositories upon peer-reviewed publication of this manuscript.

## Supplementary Tables and Figures

**Table S1. Gene usage of E1-dependent HCABs, relates to Figure 1**

IMGT gene annotations for the eleven E1-dependent HCABs by IgH (gray rows) and IgL (white rows) chains. V Germline identity represents the percent nucleotide homology to the IMGT reference of that V-gene.

**Table S2-S9. Comparison of antibody-antigen bonds in mAbClust-selected AF3 predictions and PDB experimental structures, relates to Figure 5**

HBPlus software was used to analyze the hydrogen bonds formed between antibody and antigen chains. Matched bonding pairs between AF3 predictions and experimental structures were defined as the same two residues interacting, based on alignment position, regardless of the specific atoms involved in the bond, to allow for sequence diversity between E1E2 antigens in AF3 and experimental structures. Rows are arranged in order of H77 position. Each contact is listed in the format [Protein Chain]:[Amino Acid] [Residue Number][Atoms]. The residue number is given in the Kabat scheme for antibodies, and H77 numbering for E1E2.

**Table S10. Comparison of antibody-antigen bonds in HEPC3-E2 experimental structures (PDB 6MEI and 6MEK), relates to Figure 5**

HBPlus software was used to analyze the hydrogen bonds formed between antibody and antigen chains of HEPC3-E2 PDBs 6MEK and 6MEI. Matched bonding pairs between structures were defined as the same two residues interacting, based on alignment position, regardless of the specific atoms involved in the bond, to allow for sequence diversity between the E2 antigens in PDB 6MEI (strain 1b09) and PDB 6MEK (strain 1a53). Rows are arranged in order of H77 position. Each contact is listed in the format [Protein Chain]:[Amino Acid] [Residue Number][Atoms]. The residue number is given in the Kabat scheme for antibodies, and H77 numbering for E1E2.

**Table S11. Root mean square deviation (RMSD) between CDR loops of mAbClust-selected AF3 Fv-E1E2 predictions and their corresponding experimental structures, with structures aligned by the framework regions of each Fv, relates to Figure 5.**

**Table S12. HCAB 104 Fv-E1E2 interface of the mAbClust-selected AF3 prediction, analyzed using HBPlus, relates to Figure 6**

**Table S13. Hydrogen bonds between HCAB 104 Fv-E2 and AR4A Fv-E2 (PDB 7T6X) (top) or HCAB 104 Fv-E1 and IGH505 Fv-E1 (PDB 7T6X) (bottom), relates to Figure 6**

Analysis using HBPlus. The residue number is given in the Kabat scheme for antibodies, and H77 numbering for E1E2. Rows are shaded if the two mAbs are predicted to form bonds with the same E1E2 residue.

**Fig S1. Neutralizing breadth of C117, relates to Figure 1**

Neutralizing breadth of plasma from BBAASH participant C117, who spontaneously cleared their infection, measured with a panel of antigenically diverse HCV pseudoparticles with four tiers of increasing neutralization resistance. Heatmap represents % neutralization values measured with 1:100 diluted plasma. Values are the average of two independent experiments, performed in duplicate.

**Fig S2. Binding potency of E1-dependent HCABs and reference mAbs to Bole1a E1E2, relates to Figure 1**

Binding of serial dilutions of HCABs and E1-independent (green curves) or E1-dependent (blue curves) reference HCV mAbs to lysates of Bole1a E1E2-transfected cells. HCAB curves are colored black, except for HCAB105 (burnt orange) and HCAB109 (purple) which showed low affinity binding. Data is the result of one experiment, measured in duplicate. mAbs were serially diluted five-fold from a starting concentration of 100 µg/ml. Curves were fit with a four-parameter log model and EC_50_ values calculated in GraphPad Prism. Error bars represent standard deviations.

**Fig S3. Binding competition of HEPC111 and AR5A, relates to Figure 1**

E1E2-binding competition of HEPC111 with E1-dependent reference mAbs AR4A and AR5A. Binding of biotinylated HEPC111 (black curves) to lysates of cells transfected with Bole1a E1E2 was assessed in the presence and absence of each non-biotinylated blocking antibody (HEPC111, purple; AR4A, red; AR5A, cyan). Curves were fit with a four-parameter log model in GraphPad Prism with the exception of the AR5A competition curve which was fit with a linear model due to fewer serial dilutions.

**Fig S4. Pymol alignments of mAbClust-selected AF3 Fv-E1E2 predictions and corresponding experimental structures, with structures aligned on E1 residues 314-324 (IgH505) or E2 (remaining mAbs), relates to Figure 5**

Structures were mutually trimmed to those residues that existed in both structures, using E2 for all mAbs except IGH505, which was trimmed with E1 alone. E2 or E1 from the experimental structure is shown as a transparent surface representation in gray. Fvs are colored dark red for the experimental structure, and cyan for the AF3 structure.

## References

1. Organization, W.H. (2024). Global hepatitis report 2024: action for access in low-and middle-income countries (World Health Organization).

2. Kinchen, V.J., Massaccesi, G., Flyak, A.I., Mankowski, M.C., Colbert, M.D., Osburn, W.O., Ray, S.C., Cox, A.L., Crowe, J.E., Jr., and Bailey, J.R. (2019). Plasma deconvolution identifies broadly neutralizing antibodies associated with hepatitis C virus clearance. J. Clin. Invest. 130. 10.1172/JCI130720.

3. Osburn, W.O., Snider, A.E., Wells, B.L., Latanich, R., Bailey, J.R., Thomas, D.L., Cox, A.L., and Ray, S.C. (2014). Clearance of hepatitis C infection is associated with the early appearance of broad neutralizing antibody responses. Hepatology 59, 2140–2151. 10.1002/hep.27013.

4. Park, S.B., Zimmer-Harwood, P., and Liang, T.J. (2025). Targets of protective immunity and opportunities in hepatitis C virus vaccine development. Nature Reviews Immunology. 10.1038/s41577-025-01215-9.

5. Raghuraman, S., Park, H., Osburn, W.O., Winkelstein, E., Edlin, B.R., and Rehermann, B. (2012). Spontaneous clearance of chronic hepatitis C virus infection is associated with appearance of neutralizing antibodies and reversal of T-cell exhaustion. J.Infect.Dis. 205, 763–771. jir835 [pii];10.1093/infdis/jir835 [doi].

6. Skinner, N.E., Ogega, C.O., Frumento, N., Clark, K.E., Paul, H., Yegnasubramanian, S., Schuebel, K., Meyers, J., Gupta, A., Wheelan, S., et al. (2023). Corrigendum: Convergent antibody responses are associated with broad neutralization of hepatitis C virus. Front. Immunol. 14. 10.3389/fimmu.2023.1201033.

7. Frumento, N., Figueroa, A., Wang, T., Zahid, M.N., Wang, S., Massaccesi, G., Stavrakis, G., Crowe, J.E., Jr., Flyak, A.I., Ji, H., et al. (2022). Repeated exposure to heterologous hepatitis C viruses associates with enhanced neutralizing antibody breadth and potency. J. Clin. Invest. 10.1172/JCI160058.

8. Nishio, A., Hasan, S., Park, H., Park, N., Salas, J.H., Salinas, E., Kardava, L., Juneau, P., Frumento, N., Massaccesi, G., et al. (2022). Serum neutralization activity declines but memory B cells persist after cure of chronic hepatitis C. Nature Communications 13. 10.1038/s41467-022-33035-z.

9. Skinner, N.E., and Bailey, J.R. (2020). Broadly neutralizing antibodies against hepatitis C virus: location, location, location. J. Hepatol. 72, 604–606. 10.1016/j.jhep.2020.01.005.

10. Wu, B.-R., Eltahla, A.A., Keoshkerian, E., Walker, M.R., Underwood, A., Brasher, N.A., Agapiou, D., Lloyd, A.R., and Bull, R.A. (2019). A method for detecting hepatitis C envelope specific memory B cells from multiple genotypes using cocktail E2 tetramers. J. Immunol. Methods 472, 65–74. 10.1016/j.jim.2019.06.016.

11. Ogega, C.O., Skinner, N.E., Flyak, A.I., Clark, K.E., Board, N.L., Bjorkman, P.J., Crowe, J.E., Jr., Cox, A.L., Ray, S.C., and Bailey, J.R. (2022). B cell overexpression of FCRL5 and PD-1 is associated with low antibody titers in HCV infection. PLoS Pathog. 18, e1010179. 10.1371/journal.ppat.1010179.

12. Chan, C.H., Hadlock, K.G., Foung, S.K., and Levy, S. (2001). V(H)1-69 gene is preferentially used by hepatitis C virus-associated B cell lymphomas and by normal B cells responding to the E2 viral antigen. Blood 97, 1023–1026.

13. Bailey, J.R., Flyak, A.I., Cohen, V.J., Li, H., Wasilewski, L.N., Snider, A.E., Wang, S., Learn, G.H., Kose, N., Loerinc, L., et al. (2017). Broadly neutralizing antibodies with few somatic mutations and hepatitis C virus clearance. JCI Insight 2. 10.1172/jci.insight.92872.

14. Flyak, A.I., Ruiz, S.E., Salas, J., Rho, S., Bailey, J.R., and Bjorkman, P.J. (2020). An ultralong CDRH2 in HCV neutralizing antibody demonstrates structural plasticity of antibodies against E2 glycoprotein. Elife 9. 10.7554/eLife.53169.

15. Tzarum, N., Giang, E., Kong, L., He, L., Prentoe, J., Augestad, E., Hua, Y., Castillo, S., Lauer, G.M., Bukh, J., et al. (2019). Genetic and structural insights into broad neutralization of hepatitis C virus by human VH1-69 antibodies. Sci Adv 5, eaav1882. 10.1126/sciadv.aav1882.

16. Keck, Z.Y., Wang, Y., Lau, P., Lund, G., Rangarajan, S., Fauvelle, C., Liao, G.C., Holtsberg, F.W., Warfield, K.L., Aman, M.J., et al. (2016). Affinity maturation of a broadly neutralizing human monoclonal antibody that prevents acute hepatitis C virus infection in mice. Hepatology 64, 1922–1933. 10.1002/hep.28850.

17. Pestka, J.M., Zeisel, M.B., Blaser, E., Schurmann, P., Bartosch, B., Cosset, F.L., Patel, A.H., Meisel, H., Baumert, J., Viazov, S., et al. (2007). Rapid induction of virus-neutralizing antibodies and viral clearance in a single-source outbreak of hepatitis C. Proc.Natl.Acad.Sci.U.S.A 104, 6025–6030.

18. Frumento, N., Sinnis-Bourozikas, A., Paul, H.T., Stavrakis, G., Zahid, M.N., Wang, S., Ray, S.C., Flyak, A.I., Shaw, G.M., Cox, A.L., and Bailey, J.R. (2023). Neutralizing antibodies evolve to exploit vulnerable sites in the HCV envelope glycoprotein E2 and mediate spontaneous clearance of infection. Immunity. 10.1016/j.immuni.2023.12.004.

19. Kinchen, V.J., Zahid, M.N., Flyak, A.I., Soliman, M.G., Learn, G.H., Wang, S., Davidson, E., Doranz, B.J., Ray, S.C., Cox, A.L., et al. (2018). Broadly Neutralizing Antibody Mediated Clearance of Human Hepatitis C Virus Infection. Cell Host Microbe 24, 717–730 e715. 10.1016/j.chom.2018.10.012.

20. de Jong, Y.P., Dorner, M., Mommersteeg, M.C., Xiao, J.W., Balazs, A.B., Robbins, J.B., Winer, B.Y., Gerges, S., Vega, K., Labitt, R.N., et al. (2014). Broadly neutralizing antibodies abrogate established hepatitis C virus infection. Sci. Transl. Med. 6, 254ra129. 10.1126/scitranslmed.3009512.

21. Law, M., Maruyama, T., Lewis, J., Giang, E., Tarr, A.W., Stamataki, Z., Gastaminza, P., Chisari, F.V., Jones, I.M., Fox, R.I., et al. (2008). Broadly neutralizing antibodies protect against hepatitis C virus quasispecies challenge. Nat.Med. 14, 25–27. nm1698 [pii];10.1038/nm1698 [doi].

22. Echeverría, N. (2015). Hepatitis C virus genetic variability and evolution. World J. Hepatol. 7, 831. 10.4254/wjh.v7.i6.831.

23. Salas, J.H., Urbanowicz, R.A., Guest, J.D., Frumento, N., Figueroa, A., Clark, K.E., Keck, Z., Cowton, V.M., Cole, S.J., Patel, A.H., et al. (2022). An Antigenically Diverse, Representative Panel of Envelope Glycoproteins for Hepatitis C Virus Vaccine Development. Gastroenterology 162, 562–574. 10.1053/j.gastro.2021.10.005.

24. Bailey, J.R., Laskey, S., Wasilewski, L.N., Munshaw, S., Fanning, L.J., Kenny-Walsh, E., and Ray, S.C. (2012). Constraints on viral evolution during chronic hepatitis C virus infection arising from a common-source exposure. J. Virol. 86, 12582–12590. 10.1128/JVI.01440-12.

25. Keck, Z.Y., Li, S.H., Xia, J., von Hahn, T., Balfe, P., McKeating, J.A., Witteveldt, J., Patel, A.H., Alter, H., Rice, C.M., and Foung, S.K. (2009). Mutations in hepatitis C virus E2 located outside the CD81 binding sites lead to escape from broadly neutralizing antibodies but compromise virus infectivity. J.Virol. 83, 6149–6160.

26. Bailey, J.R., Wasilewski, L.N., Snider, A.E., El-Diwany, R., Osburn, W.O., Keck, Z., Foung, S.K., and Ray, S.C. (2015). Naturally selected hepatitis C virus polymorphisms confer broad neutralizing antibody resistance. J. Clin. Invest. 125, 437–447. 10.1172/JCI78794.

27. Ogega, C.O., Skinner, N.E., Schoenle, M.V., Wilcox, X.E., Frumento, N., Wright, D.A., Paul, H.T., Sinnis-Bourozikas, A., Clark, K.E., Figueroa, A., et al. (2024). Convergent evolution and targeting of diverse E2 epitopes by human broadly neutralizing antibodies are associated with HCV clearance. Immunity. 10.1016/j.immuni.2024.03.001.

28. Keck, M.-L., Wrensch, F., Pierce, B.G., Baumert, T.F., and Foung, S.K.H. (2018). Mapping Determinants of Virus Neutralization and Viral Escape for Rational Design of a Hepatitis C Virus Vaccine. Front. Immunol. 9. 10.3389/fimmu.2018.01194.

29. Soerensen, A., Popovic, F., Olesen, C.H., Mendez, B.L., Lohse, B., Chen, Z., Farci, P., Purcell, R.H., Alter, H.J., Barfod, L.K., et al. (2025). Selection and characterization of a broadly neutralizing class of HCV anti-E2 VH1-69 antibodies. PLoS Pathog. 21, e1012428. 10.1371/journal.ppat.1012428.

30. Tzarum, N., Giang, E., Kadam, R.U., Chen, F., Nagy, K., Augestad, E.H., Velazquez-Moctezuma, R., Keck, Z.Y., Hua, Y., Stanfield, R.L., et al. (2020). An alternate conformation of HCV E2 neutralizing face as an additional vaccine target. Sci Adv 6, eabb5642. 10.1126/sciadv.abb5642.

31. Colbert, M.D., Flyak, A.I., Ogega, C.O., Kinchen, V.J., Massaccesi, G., Hernandez, M., Davidson, E., Doranz, B.J., Cox, A.L., Crowe, J.E., Jr., and Bailey, J.R. (2019). Broadly Neutralizing Antibodies Targeting New Sites of Vulnerability in Hepatitis C Virus E1E2. J. Virol. 93. 10.1128/JVI.02070-18.

32. Meunier, J.C., Russell, R.S., Goossens, V., Priem, S., Walter, H., Depla, E., Union, A., Faulk, K.N., Bukh, J., Emerson, S.U., and Purcell, R.H. (2008). Isolation and characterization of broadly neutralizing human monoclonal antibodies to the e1 glycoprotein of hepatitis C virus. J. Virol. 82, 966–973. 10.1128/JVI.01872-07.

33. Keck, Z.Y., Sung, V.M., Perkins, S., Rowe, J., Paul, S., Liang, T.J., Lai, M.M., and Foung, S.K. (2004). Human monoclonal antibody to hepatitis C virus E1 glycoprotein that blocks virus attachment and viral infectivity. J. Virol. 78, 7257–7263. 10.1128/JVI.78.13.7257-7263.2004.

34. Giang, E., Dorner, M., Prentoe, J.C., Dreux, M., Evans, M.J., Bukh, J., Rice, C.M., Ploss, A., Burton, D.R., and Law, M. (2012). Human broadly neutralizing antibodies to the envelope glycoprotein complex of hepatitis C virus. Proc.Natl.Acad.Sci.U.S.A 109, 6205–6210. 1114927109 [pii];10.1073/pnas.1114927109 [doi].

35. Torrents de la Pena, A., Sliepen, K., Eshun-Wilson, L., Newby, M.L., Allen, J.D., Zon, I., Koekkoek, S., Chumbe, A., Crispin, M., Schinkel, J., et al. (2022). Structure of the hepatitis C virus E1E2 glycoprotein complex. Science 378, 263–269. 10.1126/science.abn9884.

36. Metcalf, M.C., Janus, B.M., Yin, R., Wang, R., Guest, J.D., Pozharski, E., Law, M., Mariuzza, R.A., Toth, E.A., Pierce, B.G., et al. (2023). Structure of engineered hepatitis C virus E1E2 ectodomain in complex with neutralizing antibodies. Nat Commun 14, 3980. 10.1038/s41467-023-39659-z.

37. Abramson, J., Adler, J., Dunger, J., Evans, R., Green, T., Pritzel, A., Ronneberger, O., Willmore, L., Ballard, A.J., Bambrick, J., et al. (2024). Accurate structure prediction of biomolecular interactions with AlphaFold 3. Nature 630, 493–500. 10.1038/s41586-024-07487-w.

38. Hitawala, F.N., and Gray, J.J. (2025). What does AlphaFold3 learn about antibody and nanobody docking, and what remains unsolved? mAbs 17. 10.1080/19420862.2025.2545601.

39. Yin, R., and Pierce, B.G. (2024). Evaluation of AlphaFold antibody–antigen modeling with implications for improving predictive accuracy. Protein Sci. 33. 10.1002/pro.4865.

40. Terwilliger, T.C., Liebschner, D., Croll, T.I., Williams, C.J., McCoy, A.J., Poon, B.K., Afonine, P.V., Oeffner, R.D., Richardson, J.S., Read, R.J., and Adams, P.D. (2024). AlphaFold predictions are valuable hypotheses and accelerate but do not replace experimental structure determination. Nature Methods 21, 110–116. 10.1038/s41592-023-02087-4.

41. Cox, A.L., Netski, D.M., Mosbruger, T., Sherman, S.G., Strathdee, S., Ompad, D., Vlahov, D., Chien, D., Shyamala, V., Ray, S.C., and Thomas, D.L. (2005). Prospective evaluation of community-acquired acute-phase hepatitis C virus infection. Clin.Infect.Dis. 40, 951–958.

42. Weber, T., Potthoff, J., Bizu, S., Labuhn, M., Dold, L., Schoofs, T., Horning, M., Ercanoglu, M.S., Kreer, C., Gieselmann, L., et al. (2022). Analysis of antibodies from HCV elite neutralizers identifies genetic determinants of broad neutralization. Immunity 55, 341–354 e347. 10.1016/j.immuni.2021.12.003.

43. Guest, J.D., Wang, R., Elkholy, K.H., Chagas, A., Chao, K.L., Cleveland, T.E.t., Kim, Y.C., Keck, Z.Y., Marin, A., Yunus, A.S., et al. (2021). Design of a native-like secreted form of the hepatitis C virus E1E2 heterodimer. Proc. Natl. Acad. Sci. U. S. A. 118. 10.1073/pnas.2015149118.

44. Metcalf, M.C., Janus, B.M., Jeong, S., Law, M., Mariuzza, R.A., Toth, E.A., Pierce, B.G., Fuerst, T.R., and Ofek, G. (2025). Structure of HCV E1E2 complex reveals intersection of the homodimer interface by neutralizing antibodies. Submitted.

45. Pierce, B.G., Felbinger, N., Metcalf, M., Toth, E.A., Ofek, G., and Fuerst, T.R. (2024). Hepatitis C Virus E1E2 Structure, Diversity, and Implications for Vaccine Development. Viruses 16. 10.3390/v16050803.

46. Chen, I., Howarth, M., Lin, W., and Ting, A.Y. (2005). Site-specific labeling of cell surface proteins with biophysical probes using biotin ligase. Nature Methods 2, 99–104. 10.1038/nmeth735.

47. Huang, J., Doria-Rose, N.A., Longo, N.S., Laub, L., Lin, C.L., Turk, E., Kang, B.H., Migueles, S.A., Bailer, R.T., Mascola, J.R., and Connors, M. (2013). Isolation of human monoclonal antibodies from peripheral blood B cells. Nat. Protoc. 8, 1907–1915. 10.1038/nprot.2013.117.

48. Broering, T.J., Garrity, K.A., Boatright, N.K., Sloan, S.E., Sandor, F., Thomas, W.D., Jr., Szabo, G., Finberg, R.W., Ambrosino, D.M., and Babcock, G.J. (2009). Identification and characterization of broadly neutralizing human monoclonal antibodies directed against the E2 envelope glycoprotein of hepatitis C virus. J.Virol. 83, 12473–12482. JVI.01138-09 [pii];10.1128/JVI.01138-09 [doi].

49. Kong, L., Kadam, R.U., Giang, E., Ruwona, T.B., Nieusma, T., Culhane, J.C., Stanfield, R.L., Dawson, P.E., Wilson, I.A., and Law, M. (2015). Structure of Hepatitis C Virus Envelope Glycoprotein E1 Antigenic Site 314-324 in Complex with Antibody IGH526. J. Mol. Biol. 427, 2617–2628. 10.1016/j.jmb.2015.06.012.

50. Munshaw, S., Bailey, J.R., Liu, L., Osburn, W.O., Burke, K.P., Cox, A.L., and Ray, S.C. (2012). Computational reconstruction of Bole1a, a representative synthetic hepatitis C virus subtype 1a genome. J. Virol. 86, 5915–5921. 10.1128/JVI.05959-11.

51. Ma, Y., and Li, J. (2011). Vesicular Stomatitis Virus as a Vector To Deliver Virus-Like Particles of Human Norovirus: a New Vaccine Candidate against an Important Noncultivable Virus. J. Virol. 85, 2942–2952. 10.1128/jvi.02332-10.

52. Flyak, A.I., Ruiz, S., Colbert, M.D., Luong, T., Crowe, J.E., Jr., Bailey, J.R., and Bjorkman, P.J. (2018). HCV Broadly Neutralizing Antibodies Use a CDRH3 Disulfide Motif to Recognize an E2 Glycoprotein Site that Can Be Targeted for Vaccine Design. Cell Host Microbe 24, 703–716 e703. 10.1016/j.chom.2018.10.009.

53. Giang, E., Dorner, M., Prentoe, J.C., Dreux, M., Evans, M.J., Bukh, J., Rice, C.M., Ploss, A., Burton, D.R., and Law, M. (2012). Human broadly neutralizing antibodies to the envelope glycoprotein complex of hepatitis C virus. Proc. Natl. Acad. Sci. U. S. A. 109, 6205–6210. 10.1073/pnas.1114927109.

54. Pfaff-Kilgore, J.M., Davidson, E., Kadash-Edmondson, K., Hernandez, M., Rosenberg, E., Chambers, R., Castelli, M., Clementi, N., Mancini, N., Bailey, J.R., et al. (2022). Sites of vulnerability in HCV E1E2 identified by comprehensive functional screening. Cell Reports 39, 110859. 10.1016/j.celrep.2022.110859.

55. Dowd, K.A., Netski, D.M., Wang, X.H., Cox, A.L., and Ray, S.C. (2009). Selection Pressure from Neutralizing Antibodies Drives Sequence Evolution during Acute Infection with Hepatitis C Virus. Gastroenterology 136, 2377–2386.

56. Clipman, S.J., Mehta, S.H., Rodgers, M.A., Duggal, P., Srikrishnan, A.K., Saravanan, S., Balakrishnan, P., Vasudevan, C.K., Ray, S.C., Kumar, M.S., et al. (2021). Spatiotemporal Phylodynamics of Hepatitis C Among People Who Inject Drugs in India. Hepatology 74, 1782–1794. 10.1002/hep.31912.

57. Hedskog, C., Parhy, B., Chang, S., Zeuzem, S., Moreno, C., Shafran, S.D., Borgia, S.M., Asselah, T., Alric, L., Abergel, A., et al. (2019). Identification of 19 Novel Hepatitis C Virus Subtypes-Further Expanding HCV Classification. Open Forum Infect Dis 6, ofz076. 10.1093/ofid/ofz076.

58. Wasilewski, L.N., Ray, S.C., and Bailey, J.R. (2016). Hepatitis C virus resistance to broadly neutralizing antibodies measured using replication-competent virus and pseudoparticles. J. Gen. Virol. 97, 2883–2893. 10.1099/jgv.0.000608.

59. Prentoe, J., Velazquez-Moctezuma, R., Foung, S.K., Law, M., and Bukh, J. (2016). Hypervariable region 1 shielding of hepatitis C virus is a main contributor to genotypic differences in neutralization sensitivity. Hepatology 64, 1881–1892. 10.1002/hep.28705.

60. Stoddard, M.B., Li, H., Wang, S., Saeed, M., Andrus, L., Ding, W., Jiang, X., Learn, G.H., von Schaewen, M., Wen, J., et al. (2015). Identification, molecular cloning, and analysis of full-length hepatitis C virus transmitted/founder genotypes 1, 3, and 4. MBio 6, e02518. 10.1128/mBio.02518-14.

61. Messina, J.P., Humphreys, I., Flaxman, A., Brown, A., Cooke, G.S., Pybus, O.G., and Barnes, E. (2015). Global distribution and prevalence of hepatitis C virus genotypes. Hepatology 61, 77–87. 10.1002/hep.27259.

62. Mirabello, C., and Wallner, B. (2024/10/01). DockQ v2: improved automatic quality measure for protein multimers, nucleic acids, and small molecules. Bioinformatics 40. 10.1093/bioinformatics/btae586.

63. Basu, S., and Wallner, B. (2016). DockQ: A Quality Measure for Protein-Protein Docking Models. PLoS One 11, e0161879. 10.1371/journal.pone.0161879.

64. Martin Ester, H.-P.K., Jörg Sander, Xiaowei Xu (1996). A density-based algorithm for discovering clusters in large spatial databases with noise | Proceedings of the Second International Conference on Knowledge Discovery and Data Mining. Proceedings of the Second International Conference on Knowledge Discovery and Data Mining, 226–231. 10.5555/3001460.3001507.

65. Chen, F., Nguyen, Y.T.K., Lee, Y.-Z., Giang, E., Lau, S.C., Koide, Y., Hung, S.-H., Ueno, L., He, L., Fuerst, T.R., et al. (2025). The conserved bridging domain on HCV E1E2 glycoprotein complex is targeted by neutralizing antibodies from diverse lineages. Cold Spring Harbor Laboratory.

66. Tiller, T., Meffre, E., Yurasov, S., Tsuiji, M., Nussenzweig, M.C., and Wardemann, H. (2008). Efficient generation of monoclonal antibodies from single human B cells by single cell RT-PCR and expression vector cloning. J. Immunol. Methods 329, 112–124. 10.1016/j.jim.2007.09.017.

67. Bailey, J.R., Urbanowicz, R.A., Ball, J.K., Law, M., and Foung, S.K.H. (2019). Standardized Method for the Study of Antibody Neutralization of HCV Pseudoparticles (HCVpp). Methods Mol. Biol. 1911, 441–450. 10.1007/978-1-4939-8976-8_30.

68. Kuiken, C., Yusim, K., Boykin, L., and Richardson, R. (2005). The Los Alamos hepatitis C sequence database. Bioinformatics 21, 379–384.

69. Combet, C., Garnier, N., Charavay, C., Grando, D., Crisan, D., Lopez, J., Dehne-Garcia, A., Geourjon, C., Bettler, E., Hulo, C., et al. (2007). euHCVdb: the European hepatitis C virus database. Nucleic Acids Res. 35, D363–D366. 10.1093/nar/gkl970.

70. McDonald, I.K., and Thornton, J.M. (1994). Satisfying Hydrogen Bonding Potential in Proteins. J. Mol. Biol. 238, 777–793. 10.1006/jmbi.1994.1334.

71. Krissinel, E., and Henrick, K. (2007). Inference of Macromolecular Assemblies from Crystalline State. J. Mol. Biol. 372, 774–797. 10.1016/j.jmb.2007.05.022.

72. Tange, O. (2018). GNU Parallel (Ole Tange). 10.5281/zenodo.1146014.

