## Supplementary Materials for "mAbClust with AlphaFold 3 avoids hallucinations to define a quaternary broadly neutralizing HCV epitope"

**Table S1**

| HCAB | V-gene | V Germline Identity (%) | D-gene | J-gene | CDR3 Length |
|---|---|---|---|---|---|
| 100 | IGHV3-30-3 | 92.01 | IGHD3-9 | IGHJ6 | 25 |
| | IGKV4-1 | 97.57 | | IGKJ4 | 11 |
| 102 | IGHV5-51 | 93.75 | IGHD2-15 | IGHJ2 | 20 |
| | IGLV1-51 | 96.79 | | IGLJ2 | 13 |
| 103 | IGHV1-69 | 87.85 | IGHD2-2 | IGHJ4 | 14 |
| | IGKV3-20 | 94.92 | | IGKJ1 | 10 |
| 104 | IGHV4-4 | 95.09 | IGHD3-10 | IGHJ6 | 22 |
| | IGKV1-12 | 95.65 | | IGKJ4 | 11 |
| 105 | IGHV1-24 | 93.75 | IGHD1-26 | IGHJ4 | 17 |
| | IGKV1-39 | 100 | | IGKJ1 | 11 |
| 108 | IGHV1-69 | 89.93 | IGHD5-18 | IGHJ5 | 19 |
| | IGKV1-39 | 92.49 | | IGKJ1 | 11 |
| 109 | IGHV3-30-3 | 93.75 | IGHD5-12 | IGHJ6 | 19 |
| | IGLV3-9 | 95.99 | | IGLJ1 | 11 |
| 110 | IGHV3-30-3 | 89.24 | IGHD6-19 | IGHJ5 | 16 |
| | IGKV4-1 | 95.57 | | IGKJ1 | 11 |
| 111 | IGHV3-30 | 92.01 | IGHD2-2 | IGHJ4 | 17 |
| | IGKV2-28 | 95.9 | | IGKJ1 | 11 |
| 114 | IGHV4-39 | 96.43 | IGHD1-7 | IGHJ4 | 21 |
| | IGKV1-16 | 96.84 | | IGKJ5 | 11 |
| 115 | IGHV5-51 | 92.71 | IGHD2-15 | IGHJ6 | 19 |
| | IGKV1-5 | 94.07 | | IGKJ1 | 12 |

**Table S1. Gene usage of E1-dependent HCABs, relates to Figure 1**

IMGT gene annotations for the eleven E1-dependent HCABs by IgH (gray rows) and IgL (white rows) chains. V Germline identity represents the percent nucleotide homology to the IMGT reference of that V-gene.

### Table S2

| HEPC74 AF3 | | | HEPC74 PDB 6MEH | | |
|---|---|---|---|---|---|
| Antibody Contact | Distance | Antigen Contact | Antibody Contact | Distance | Antigen Contact |
| | | | H:GLY H100C[O] | 2.69 Å | E2:HIS 421[NE2] |
| H:CYS H100[N, O] | 2.90 Å | E2:CYS 429[N, O] | | | |
| H:LYS H98[O] | 2.81 Å | E2:GLU 431[N] | H:LYS H98[O] | 3.04 Å | E2:ASP 431[N] |
| | | | L:THR L56[OG1] | 3.09 Å | E2:SER 432[O] |
| H:ARG H94[NH1, NH2] | 2.77-3.13 Å | E2:ASN 434[O] | H:ARG H94[NH1] | 2.97 Å | E2:HIS 434[O] |
| H:LYS H98[NZ] | 2.87 Å | E2:ASN 434[OD1] | | | |
| L:GLU L55[OE1] | 2.24 Å | E2:ASN 434[ND2] | L:GLU L55[OE1] | 2.93 Å | E2:HIS 434[NE2] |
| | | | H:TYR H32[OH] | 2.81 Å | E2:THR 435[O] |
| H:ILE H30[N] | 2.85 Å | E2:LYS 446[O] | H:ILE H30[N] | 2.88 Å | E2:LYS 446[O] |
| H:GLY H27[O] | 3.28 Å | E2:ASN 448[ND2] | H:GLY H27[O] | 2.98 Å | E2:ASN 448[ND2] |
| H:THR H28[O] | 2.83 Å | E2:ASN 448[N] | H:THR H28[O] | 2.93 Å | E2:ASN 448[N] |
| | | | H:GLY H100B[N] | 2.24 Å | E2:GLU 531[OE2] |

### Table S3

| AR3A AF3 | | | AR3A PDB 6BKB | | |
|---|---|---|---|---|---|
| Antibody Contact | Distance | Antigen Contact | Antibody Contact | Distance | Antigen Contact |
| H:CYS H100A[N, O] | 2.83-3.02 Å | E2:CYS 429[N, O] | H:CYS H100A[N, O] | 2.82-3.27 Å | E2:CYS 429[N, O] |
| L:TYR L32[OH] | 2.58 Å | E2:ASN 430[OD1] | L:TYR L32[OH] | 2.62 Å | E2:ASN 430[OD1] |
| H:ARG H99[NH1, O] | 2.87-3.27 Å | E2:ASP 431[N, OD2] | H:ARG H99[NH1, O] | 2.98-3.30 Å | E2:ASP 431[N, OD2] |
| H:GLU H96[O] | 2.83 Å | E2:TYR 443[OH] | H:GLU H96[O] | 2.28 Å | E2:TYR 443[OH] |
| | | | H:SER H100B[OG] | 3.30 Å | E2:GLU 531[OE2] |

### Table S4

| HEPC3 AF3 | | | HEPC3 PDB 6MEK | | |
|---|---|---|---|---|---|
| Antibody Contact | Distance | Antigen Contact | Antibody Contact | Distance | Antigen Contact |
| H:CYS H100[N, O] | 2.91-3.01 Å | E2:CYS 429[N, O] | H:CYS H100[N, O] | 2.95-3.09 Å | E2:CYS 429[N, O] |
| H:ARG H98[NE, NH2, O] | 2.83-3.29 Å | E2:ASP 431[N, O] | H:ARG H98[NE, O] | 2.82 Å | E2:ASP 431[N, O] |
| | | | H:GLN H1[N] | 3.02 Å | E2:LEU 433[O] |
| | | | H:GLN H1[N] | 3.09 Å | E2:ASN 434[O] |
| H:ASP H101[O] | 2.79 Å | E2:ASN 434[ND2] | | | |
| H:VAL H2[N] | 2.90 Å | E2:ASN 434[O] | H:VAL H2[N] | 3.03 Å | E2:ASN 434[O] |
| H:ARG H94[NH1] | 2.16 Å | E2:ASN 434[OD1] | H:ARG H94[NH1] | 2.31 Å | E2:ASN 434[OD1] |
| H:THR H28[N, OG1] | 2.79-2.96 Å | E2:THR 435[O] | H:THR H28[N] | 2.99 Å | E2:THR 435[O] |
| H:GLU H33[OE1] | 2.39 Å | E2:TYR 443[OH] | | | |
| H:ASN H30[N] | 2.78 Å | E2:LYS 446[O] | H:ASN H30[N] | 2.82 Å | E2:LYS 446[O] |
| H:GLU H73[OE2] | 2.63 Å | E2:LYS 446[NZ] | | | |
| H:THR H28[O] | 2.89 Å | E2:ASN 448[N] | H:THR H28[O] | 2.74 Å | E2:ASP 448[N] |

### Table S5

| HCAB64 AF3 | | | HCAB64 PDB 8W0W | | |
|---|---|---|---|---|---|
| Antibody Contact | Distance | Antigen Contact | Antibody Contact | Distance | Antigen Contact |
| H:CYS H100[N, O] | 3.01-3.18 Å | E2:CYS 429[N, O] | H:CYS H100[N] | 3.17 Å | E2:CYS 429[O] |
| | | | H:ARG H96[NH1] | 3.00 Å | E2:ASP 431[OD2] |
| H:GLY H98[O] | 2.92 Å | E2:GLU 431[N] | H:GLY H98[O] | 2.88 Å | E2:ASP 431[N] |
| H:VAL H2[N] | 2.66 Å | E2:ASN 434[O] | H:VAL H2[N] | 2.91 Å | E2:HIS 434[O] |
| H:ARG H96[NH2] | 3.23 Å | E2:ASN 434[OD1] | | | |
| L:THR L56[OG1] | 3.28 Å | E2:ASN 434[OD1] | | | |
| H:THR H28[N, OG1] | 2.76-2.89 Å | E2:THR 435[O] | H:THR H28[N] | 2.81 Å | E2:THR 435[O] |
| | | | H:TYR H29[N] | 3.36 Å | E2:THR 435[O] |
| H:ILE H30[N] | 2.91 Å | E2:LYS 446[O] | H:ILE H30[N] | 3.21 Å | E2:LYS 446[O] |
| H:THR H52A[OG1] | 2.73 Å | E2:LYS 446[NZ] | | | |
| H:ASP H53[OD1] | 2.28 Å | E2:LYS 446[NZ] | H:ASP H53[OD1] | 2.99 Å | E2:LYS 446[NZ] |
| | | | H:TYR H27[O] | 2.88 Å | E2:ASN 448[ND2] |
| H:THR H28[O] | 2.91 Å | E2:ASN 448[N] | H:THR H28[O] | 2.76 Å | E2:ASN 448[N] |
| | | | H:ARG H100B[NE] | 2.01 Å | E2:GLU 531[OE2] |

### Table S6

| HCAB55 AF3 | | | HCAB55 PDB 8W0V | | |
|---|---|---|---|---|---|
| **Antibody Contact** | **Distance** | **Antigen Contact** | **Antibody Contact** | **Distance** | **Antigen Contact** |
| H:CYS H100[N, O] | 2.91-3.13 Å | E2:CYS 429[N, O] | H:CYS H100[N] | 3.09 Å | E2:CYS 429[O] |
| H:ARG H96[NH1] | 2.79 Å | E2:ASP 431[OD2] | H:ARG H96[NH1] | 3.20 Å | E2:ASP 431[OD2] |
| H:ALA H98[O] | 2.81 Å | E2:ASP 431[N] | H:ALA H98[O] | 2.77 Å | E2:ASP 431[N] |
| H:VAL H2[N] | 2.89 Å | E2:GLN 434[O] | H:VAL H2[N] | 3.06 Å | E2:HIS 434[O] |
| H:THR H28[N, OG1] | 2.72-3.19 Å | E2:THR 435[O] | H:THR H28[N] | 2.89 Å | E2:THR 435[O] |
| | | | H:PHE H29[N] | 3.21 Å | E2:THR 435[O] |
| H:ILE H30[N] | 2.99 Å | E2:LYS 446[O] | H:ILE H30[N] | 2.87 Å | E2:LYS 446[O] |
| | | | H:TYR H27[O] | 3.05 Å | E2:ASN 448[ND2] |
| H:THR H28[O] | 2.90 Å | E2:ASN 448[N] | H:THR H28[O] | 2.92 Å | E2:ASN 448[N] |
| | | | H:ARG H100A[NH1] | 2.63 Å | E2:GLU 531[OE1] |

### Table S7

| HEPC46 AF3 | | | HEPC46 PDB 6MEK | | |
|---|---|---|---|---|---|
| **Antibody Contact** | **Distance** | **Antigen Contact** | **Antibody Contact** | **Distance** | **Antigen Contact** |
| H:SER H31[O] | 2.48 Å | E2:THR 542[OG1] | H:SER H31[O] | 2.93 Å | E2:THR 542[OG1] |
| H:GLN H97[OE1] | 3.13 Å | E2:ARG 543[NH1] | H:GLN H97[OE1] | 3.33 Å | E2:ARG 543[NE] |
| H:SER H52[OG] | 3.39 Å | E2:LEU 546[O] | H:SER H52[OG] | 2.35 Å | E2:LEU 546[O] |
| L:SER L30[O] | 2.74 Å | E2:ARG 596[NH2] | | | |

### Table S8

| HCAB40 AF3 | | | HCAB40 PDB 8W0X | | |
|---|---|---|---|---|---|
| **Antibody Contact** | **Distance** | **Antigen Contact** | **Antibody Contact** | **Distance** | **Antigen Contact** |
| | | | H:SER H74[O] | 3.26 Å | E2:HIS 445[NE2] |
| | | | H:SER H74[O] | 2.39 Å | E2:LYS 446[NZ] |
| H:ASP H100E[OD2] | 2.48 Å | E2:ARG 587[NH2] | | | |
| | | | H:SER H30[OG] | 2.50 Å | E2:VAL 622[O] |
| H:ARG H73[NH1, NH2] | 2.78-3.16 Å | E2:ASN 623[OD1] | H:ARG H73[NH1] | 2.90 Å | E2:ASN 623[OD1] |
| H:GLY H31[O] | 2.81 Å | E2:ILE 626[N] | H:GLY H31[O] | 2.80 Å | E2:ILE 626[N] |
| H:TYR H100[N] | 2.89 Å | E2:LYS 628[O] | H:TYR H100[N] | 3.08 Å | E2:LYS 628[O] |
| H:PRO H98[O] | 2.86 Å | E2:LYS 628[N] | H:PRO H98[O] | 3.00 Å | E2:LYS 628[N] |

### Table S9

| IGH505 AF3 | | | IGH505 PDB 7T6X | | |
|---|---|---|---|---|---|
| **Antibody Contact** | **Distance** | **Antigen Contact** | **Antibody Contact** | **Distance** | **Antigen Contact** |
| H:TYR H98[OH] | 2.62 Å | E1:HIS 316[ND1] | H:TYR H98[OH] | 2.98 Å | E1:HIS 316[ND1] |
| L:ASN L31[OD1] | 3.01 Å | E1:HIS 316[N] | L:ASN L31[OD1] | 3.04 Å | E1:HIS 316[N] |
| H:ASP H95[OD1] | 3.18 Å | E1:TRP 320[NE1] | H:ASP H95[OD1] | 3.03 Å | E1:TRP 320[NE1] |
| H:TYR H100B[OH] | 2.80 Å | E1:ASP 321[OD1] | | | |
| H:ASN H52[ND2] | 3.29 Å | E1:MET 323[O] | H:ASN H52[ND2] | 2.90 Å | E1:MET 323[O] |
| | | | H:ASN H56[ND2] | 3.12 Å | E1:MET 323[O] |
| | | | H:ASN H52[ND2] | 2.99 Å | E1:MET 324[O] |
| H:HIS H100[NE2] | 3.37 Å | E1:ARG 339[NE] | | | |

**Table S2-S9. Comparison of antibody-antigen bonds in mAbClust-selected AF3 predictions and PDB experimental structures, relates to Figure 5**

HBPlus software was used to analyze the hydrogen bonds formed between antibody and antigen chains. Matched bonding pairs between AF3 predictions and experimental structures were defined as the same two residues interacting, based on alignment position, regardless of the specific atoms involved in the bond, to allow for sequence diversity between E1E2 antigens in AF3 and experimental structures. Rows are arranged in order of H77 position. Each contact is listed in the format [Protein Chain]:[Amino Acid] [Residue Number][Atoms]. The residue number is given in the Kabat scheme for antibodies, and H77 numbering for E1E2.

### Table S10

| HEPC3 PDB 6MEK | | | HEPC3 PDB 6MEI | | |
|---|---|---|---|---|---|
| **Antibody Contact** | **Distance** | **Antigen Contact** | **Antibody Contact** | **Distance** | **Antigen Contact** |
| H:CYS H100[N, O] | 2.95-3.09 Å | E2:CYS 429[N, O] | H:CYS H100[N, O] | 2.89-3.12 Å | E2:CYS 429[N, O] |
| H:ARG H98[NE, O] | 2.82 Å | E2:ASP 431[N, O] | H:ARG H98[O] | 2.97 Å | E2:ASP 431[N] |
| H:GLN H1[N] | 3.02 Å | E2:LEU 433[O] | | | |
| H:GLN H1[N] | 3.09 Å | E2:ASN 434[O] | | | |
| | | | H:ASP H101[OD2] | 2.94 Å | E2:HIS 434[NE2] |
| H:VAL H2[N] | 3.03 Å | E2:ASN 434[O] | | | |
| H:ARG H94[NH1] | 2.31 Å | E2:ASN 434[OD1] | | | |
| H:THR H28[N] | 2.99 Å | E2:THR 435[O] | H:THR H28[OG1] | 3.11 Å | E2:THR 435[O] |
| H:ASN H30[N] | 2.82 Å | E2:LYS 446[O] | H:ASN H30[N] | 2.85 Å | E2:LYS 446[O] |
| | | | H:PRO H53[O] | 2.33 Å | E2:LYS 446[NZ] |
| | | | H:GLY H27[O] | 2.93 Å | E2:ASN 448[ND2] |
| H:THR H28[O] | 2.74 Å | E2:ASP 448[N] | H:THR H28[O] | 2.84 Å | E2:ASN 448[N] |

**Table S10. Comparison of antibody-antigen bonds in HEPC3-E2 experimental structures (PDB 6MEI and 6MEK), relates to Figure 5**

HBPlus software was used to analyze the hydrogen bonds and salt bridges formed between antibody and antigen chains of HEPC3-E2 PDBs 6MEK and 6MEI. Matched bonding pairs between structures were defined as the same two residues interacting, based on alignment position, regardless of the specific atoms involved in the bond, to allow for sequence diversity between the E2 antigens in PDB 6MEI (strain 1b09) and PDB 6MEK (strain 1a53). Rows are arranged in order of H77 position. Each contact is listed in the format [Protein Chain]:[Amino Acid] [Residue Number][Atoms]. The residue number is given in the Kabat scheme for antibodies, and H77 numbering for E2.

### Table S11

| Antibody | CDRH1 | CDRH2 | CDRH3 | CDRL1 | CDRL2 | CDRL3 |
|---|---|---|---|---|---|---|
| AR3A | 0.52 | 0.8 | 0.83 | 0.9 | 0.56 | 0.56 |
| HCAB40 | 0.3 | 1.19 | 2.19 | 0.19 | 0.35 | 0.33 |
| HCAB55 | 0.97 | 1.08 | 1.9 | 0.45 | 0.37 | 0.67 |
| HCAB64 | 1.01 | 1.13 | 1.27 | 0.31 | 0.44 | 0.54 |
| HEPC3 | 0.33 | 0.49 | 0.46 | 0.3 | 0.47 | 0.57 |
| HEPC46 | 0.46 | 1.11 | 0.63 | 0.69 | 0.49 | 1.89 |
| HEPC74 | 1.88 | 0.95 | 2.14 | 0.21 | 0.26 | 2.31 |
| IGH505 | 1.08 | 0.72 | 1.77 | 0.84 | 4.94 | 0.63 |
| 6MEK vs 6MEI | 0.63 | 0.28 | 1.05 | 0.4 | 0.29 | 0.51 |

**Table S11. Root mean square deviation (RMSD) between CDR loops of mAbClust-selected AF3 Fv-E1E2 predictions and their corresponding experimental structures, with structures aligned by the framework regions of each Fv. Relates to Figure 5**

### Table S12

| Antibody Contact | Distance | Antigen Contact |
|---|---|---|
| H:SER H28[OG] | 3.14 Å | E1:ILE 313[O] |
| H:ASN H30[ND2] | 2.76 Å | E1:MET 318[SD] |
| L:ALA L91[O] | 2.53 Å | E2:ARG 648[NH1] |
| H:ASP H100H[OD1] | 2.85 Å | E2:ARG 648[NH2] |
| H:GLY H100[O] | 2.58 Å | E2:GLY 649[N] |
| H:SER H100A[N] | 2.83 Å | E2:ASN 695[O] |
| H:TYR H97[O] | 2.57 Å | E2:ASN 695[ND2] |
| H:SER H100A[O] | 2.94 Å | E2:VAL 697[N] |
| H:MET H100C[N] | 2.76 Å | E2:ASP 698[OD1] |
| H:TYR H33[OH] | 2.66 Å | E2:TYR 701[OH] |
| H:TYR H33[OH] | 2.66 Å | E2:TYR 701[OH] |

**Table S12. HCAB 104 Fv-E1E2 interface of the mAbClust-selected AF3 prediction, analyzed using HBPlus, relates to Figure 6**

### Table S13

| H77 Pos | HCAB104 AF3 Antibody Contact | HCAB104 AF3 Distance | HCAB104 AF3 Antigen Contact | AR4A PDB 7T6X Antibody Contact | AR4A PDB 7T6X Distance | AR4A PDB 7T6X Antigen Contact |
|---|---|---|---|---|---|---|
| 313 | H:SER H28[OG] | 3.14 Å | E1:ILE 313[O] | | | |
| 318 | H:ASN H30[ND2] | 2.76 Å | E1:MET 318[SD] | | | |
| 626 | | | | H:ARG H100G[NH2] | 2.84 Å | E2:ARG 626[O] |
| 648 | H:ASP H100H[OD1]<br>L:ALA L91[O] | 2.53-2.85 Å | E2:ARG 648[NH1, NH2] | | | |
| 649 | H:GLY H100[O] | 2.58 Å | E2:GLY 649[N] | H:THR H100C[O] | 3.00 Å | E2:GLY 649[N] |
| 650 | | | | H:THR H100C[OG1] | 2.76 Å | E2:GLU 650[O] |
| 695 | H:TYR H97[O]<br>H:SER H100A[N] | 2.57-2.83 Å | E2:ASN 695[ND2, O] | H:PHE H100D[N] | 3.03 Å | E2:ASN 695[O] |
| 697 | H:SER H100A[O] | 2.94 Å | E2:VAL 697[N] | H:PHE H100D[O] | 2.88 Å | E2:VAL 697[N] |
| 698 | H:MET H100C[N] | 2.76 Å | E2:ASP 698[OD1] | H:TRP H100F[N] | 2.84 Å | E2:ASP 698[OD1] |
| 701 | H:TYR H33[OH] | 2.66 Å | E2:TYR 701[OH] | | | |

| H77 Pos | HCAB104 AF3 Antibody Contact | HCAB104 AF3 Distance | HCAB104 AF3 Antigen Contact | IGH505 PDB 7T6X Antibody Contact | IGH505 PDB 7T6X Distance | IGH505 PDB 7T6X Antigen Contact |
|---|---|---|---|---|---|---|
| 313 | H:SER H28[OG] | 3.14 Å | E1:ILE 313[O] | | | |
| 316 | | | | H:TYR H98[OH]<br>L:ASN L31[OD1] | 2.98-3.04 Å | E1:HIS 316[N, ND1] |
| 318 | H:ASN H30[ND2] | 2.76 Å | E1:MET 318[SD] | | | |
| 320 | | | | H:ASP H95[OD1] | 3.03 Å | E1:TRP 320[NE1] |
| 323 | | | | H:ASN H52[ND2]<br>H:ASN H56[ND2] | 2.90-3.12 Å | E1:MET 323[O] |
| 324 | | | | H:ASN H52[ND2] | 2.99 Å | E1:MET 324[O] |
| 648 | H:ASP H100H[OD1]<br>L:ALA L91[O] | 2.53-2.85 Å | E2:ARG 648[NH1, NH2] | | | |
| 649 | H:GLY H100[O] | 2.58 Å | E2:GLY 649[N] | | | |
| 695 | H:TYR H97[O]<br>H:SER H100A[N] | 2.57-2.83 Å | E2:ASN 695[ND2, O] | | | |
| 697 | H:SER H100A[O] | 2.94 Å | E2:VAL 697[N] | | | |
| 698 | H:MET H100C[N] | 2.76 Å | E2:ASP 698[OD1] | | | |
| 701 | H:TYR H33[OH] | 2.66 Å | E2:TYR 701[OH] | | | |

**Table S13. Hydrogen bonds between HCAB 104 Fv-E2 and AR4A Fv-E2 (PDB 7T6X) (top) or HCAB 104 Fv-E1 and IGH505 Fv-E1 (PDB 7T6X) (bottom), relates to Figure 6**

Analysis using HBPlus. The residue number is given in the Kabat scheme for antibodies, and H77 numbering for E1E2. Rows are shaded if the two mAbs are predicted to form bonds with the same E1E2 residue.

### Figure S1

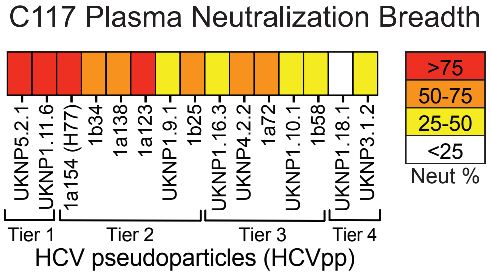


**Fig S1. Neutralizing breadth of C117, relates to Figure 1**

Neutralizing breadth of plasma from BBAASH participant C117, who spontaneously cleared their infection, measured with a panel of antigenically diverse HCV pseudoparticles with four tiers of increasing neutralization resistance. Heatmap represents % neutralization values measured with 1:100 diluted plasma. Values are the average of two independent experiments, performed in duplicate.

### Figure S2

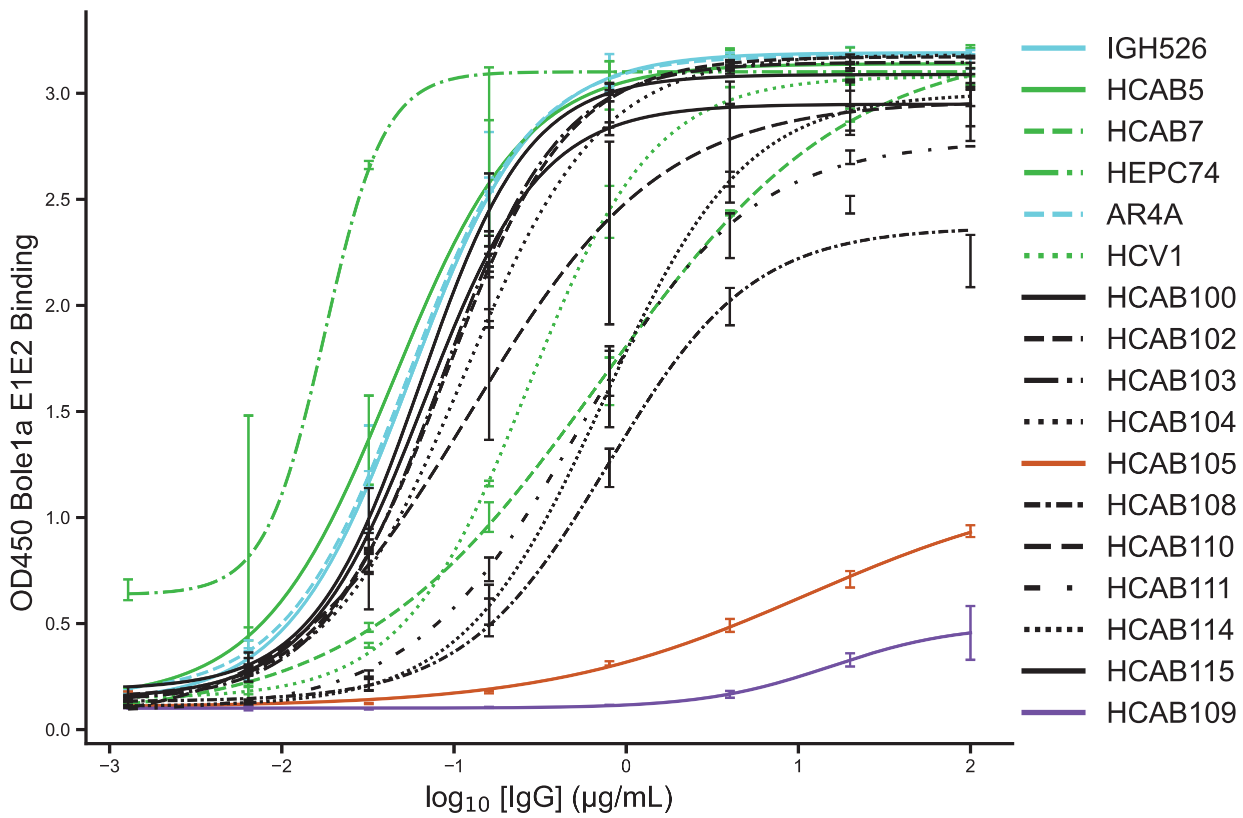


**Fig S2. Binding potency of E1-dependent HCABs and reference mAbs to Bole1a E1E2, relates to Figure 1**

Binding of serial dilutions of HCABs and E1-independent (green curves) or E1-dependent (blue curves) reference HCV mAbs to lysates of Bole1a E1E2-transfected cells. HCAB curves are colored black, except for HCAB105 (burnt orange) and HCAB109 (purple) which showed low affinity binding. Data is the result of one experiment, measured in duplicate. mAbs were serially diluted five-fold from a starting concentration of 100 μg/ml. Curves were fit with a four-parameter log model and EC50 values calculated in GraphPad Prism. Error bars represent standard deviations.

**Figure S3**

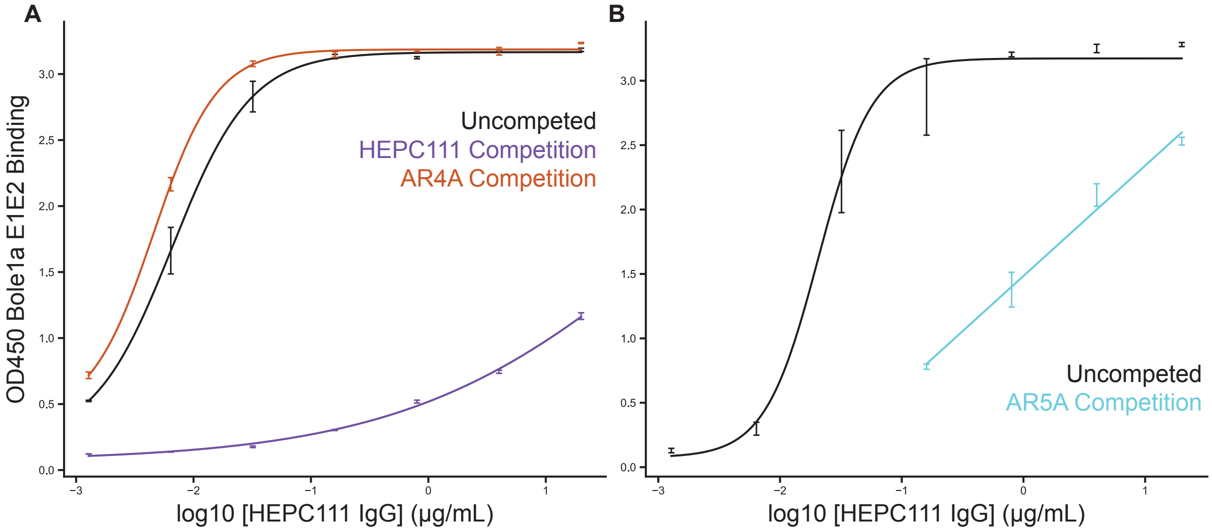


**Fig S3. Binding competition of HEPC111 and AR5A, relates to Figure 1**

E1E2-binding competition of HEPC111 with E1-dependent reference mAbs AR4A and AR5A. Binding of biotinylated HEPC111 (black curves) to lysates of cells transfected with Bole1a E1E2 was assessed in the presence and absence of each non-biotinylated blocking antibody (HEPC111, purple; AR4A, red; AR5A, cyan). Curves were fit with a four-parameter log model in GraphPad Prism with the exception of the AR5A competition curve which was fit with a linear model due to fewer serial dilutions.

**Figure S4**

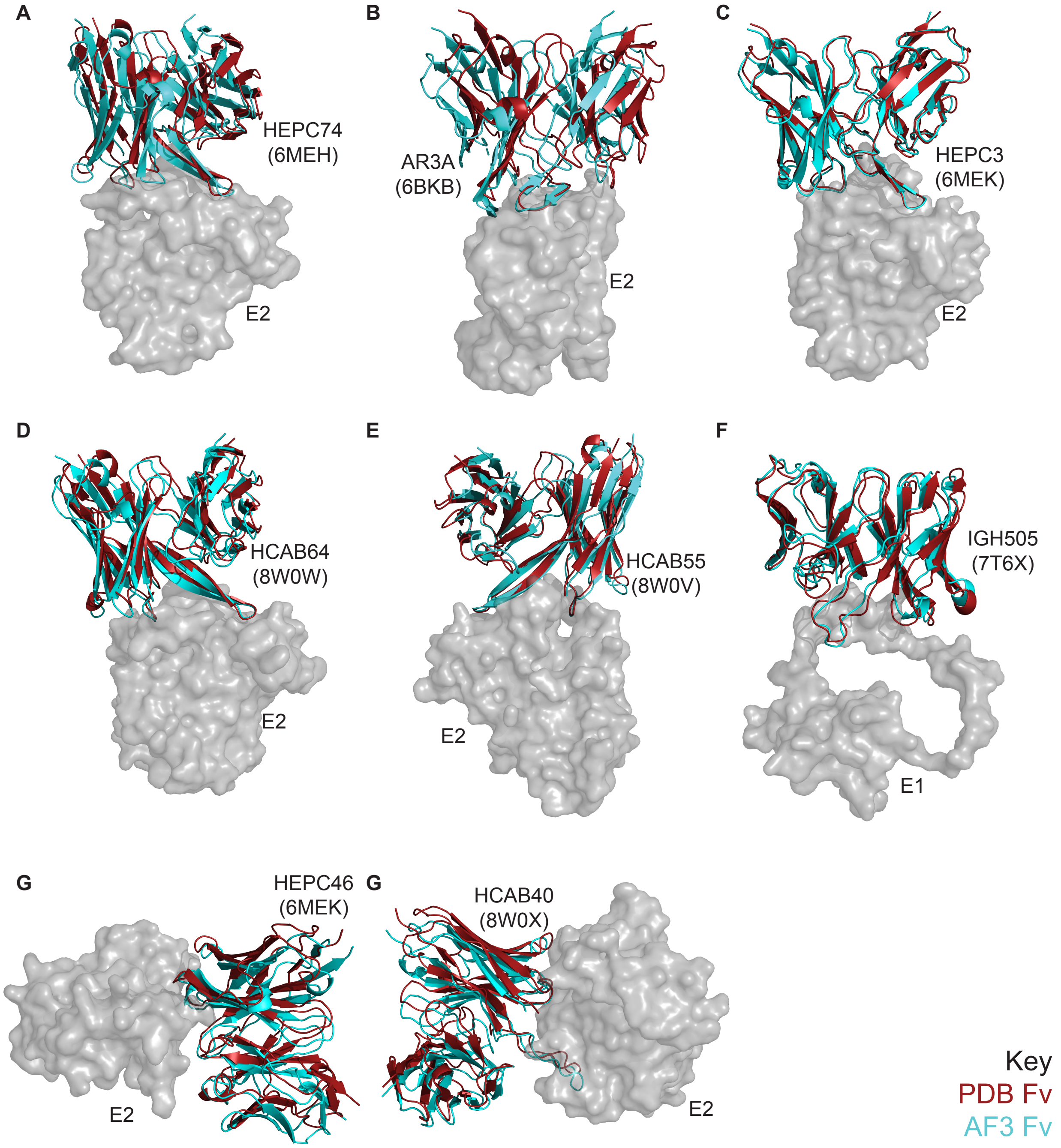


**Fig S4. Pymol alignments of mAbClust-selected AF3 Fv-E1E2 predictions and corresponding experimental structures, with structures aligned on E1 residues 314-324 (IgH505) or E2 (remaining mAbs), relates to Figure 5**

Structures were mutually trimmed to those residues that existed in both structures, using E2 for all mAbs except IGH505, which was trimmed with E1 alone. E2 or E1 from the experimental structure is shown as a transparent surface representation in gray. Fvs are colored dark red for the experimental structure, and cyan for the AF3 structure.
